# Human hepatocyte PNPLA3 148M exacerbates rapid non-alcoholic steatohepatitis development in chimeric mice

**DOI:** 10.1101/2020.11.19.387613

**Authors:** Mohammad Kabbani, Eleftherios Michailidis, Sandra Steensels, Clifton G. Fulmer, Joseph M. Luna, Jérémie Le Pen, Matteo Tardelli, Brandon Razooky, Inna Ricardo-Lax, Chenhui Zou, Briana Zeck, Ansgar F. Stenzel, Corrine Quirk, Lander Foquet, Alison W. Ashbrook, William M. Schneider, Serkan Belkaya, Gadi Lalazar, Yupu Liang, Meredith Pittman, Lindsey Devisscher, Hiroshi Suemizu, Neil D. Theise, Luis Chiriboga, David E. Cohen, Robert Copenhaver, Markus Grompe, Philip Meuleman, Baran A. Ersoy, Charles M. Rice, Ype P. de Jong

## Abstract

Advanced non-alcoholic fatty liver disease (NAFLD) is a rapidly emerging global health problem associated with pre-disposing genetic polymorphisms, most strikingly an isoleucine to methionine substitution in patatin-like phospholipase domain-containing protein 3 (PNPLA3-I148M). Here, we study how human hepatocytes with PNPLA3 148I and 148M variants engrafted in the livers of chimeric mice respond to a hypercaloric Western-style diet. As early as 4 weeks, mice developed dyslipidemia, impaired glucose tolerance, and steatohepatitis selectively in the human graft, followed by pericellular fibrosis after 8 weeks of hypercaloric feeding. The PNPLA3 148M variant, either from a homozygous 148M human donor or overexpressed in a homozygous 148I donor background, caused widespread microvesicular steatosis and even more severe steatohepatitis. We conclude that PNPLA3 148M in human hepatocytes exacerbates NAFLD. These models will facilitate mechanistic studies into human genetic variants associated with advanced fatty liver diseases.

## Introduction

Non-alcoholic fatty liver disease (NAFLD) has become the most prevalent liver disease in many countries, affecting approximately 25% of the world population.^1^ NAFLD is associated with obesity, insulin resistance, dyslipidemia and hypertension.^2^ A subset of patients with NAFLD develop non-alcoholic steatohepatitis (NASH), which is widely believed to drive fibrosis progression towards advanced liver disease including cirrhosis and hepatocellular carcinoma (HCC). ^3^

The complex nutritional and environmental factors leading to NAFLD remain to be better defined. ^4^ In addition, a number of genetic variants have been associated with advanced NAFLD.^5^ The first genetic polymorphism identified in a population with increased hepatic steatosis was rs738409 in patatin-like phospholipase domain containing protein-3 (*PNPLA3*).^6^ This polymorphism encodes for an isoleucine to methionine substitution at codon position 148 in PNPLA3. Since its identification by Hobbs and colleagues, the PNPLA3-I148M variant has been globally associated with advanced NAFLD and other liver diseases including HCC.^7^

The mechanisms by which PNPLA3-148M drives advanced NAFLD remains an active area of investigation.^8, 9^ Although PNPLA3 is expressed in hepatocytes, hepatic stellate cells and adipocytes, accumulating evidence from mouse models indicates that hepatocyte-specific PNPLA3-148M promotes the pathogenesis of NAFLD. Transgenic overexpression of human PNPLA3-148M in mouse hepatocytes but not adipocytes exacerbated hepatic steatosis.^10^ And while increased hepatic fat content was observed in *Pnpla3*-148M knock-in mice,^11^ this was reversed by silencing *Pnpla3* in hepatocytes.^12^ Functionally, PNPLA3 exhibits hydrolase activity towards triglycerides and acyltransferase activity for polyunsaturated fatty acids in phospholipids.^8^ PNPLA3-148M differs from 148I in several ways: PNPLA3-148M was shown to have decreased hydrolase activity against triglycerides,^13^ increased acyltransferase activity for lysophosphatidic acid;^14^ resistance to ubiquitylation and proteasomal degradation;^15^ and inhibition of adipose triglyceride lipase on lipid droplets via interaction with α/β-hydrolase domain containing 5.^16,17^ How these and other mechanistic differences drive NAFLD progression remains to be further elucidated.

Despite the vast disease burden there are no approved pharmacological therapies for NASH and sustained weight loss remains the only proven intervention. Given its complex pathogenesis, many different pathways are under pre-clinical and clinical investigation, some of which have resulted in compounds that progressed to Phase 3 trials.^18^ Several of these pathways were identified in murine NASH models that share limited features with human disease.^19^ Furthermore, investigational strategies that target subsets of NASH patients, e.g. those with genetic variants, are further complicated by species differences. In the case of PNPLA3, there is low sequence homology between the human and mouse orthologues. The human protein is 97 amino acids longer than the mouse PNPLA3. The relative mRNA expression starkly differs between adipose tissue and liver in each species,^20,21^ and it remains unclear if the same transcription factors regulate its expression in murine and human hepatocytes.^22,23^ Elucidation of the precise mechanisms by which human genetic variants in hepatocytes affect steatohepatitis would be aided by minimizing species-specific variances in a physiologically relevant experimental setting.

Here, we established a human hepatocyte NAFLD model. We applied this model to study human hepatocyte PNPLA3-148M, either from a 148M homozygous human hepatocyte donor and by overexpressing PNPLA3 variants in a 148I homozygous donor background. Our results show that human hepatocytes swiftly develop steatohepatitis in response to hypercaloric feeding, which is exacerbated by the presence of PNPLA3-148M in hepatocytes. These models will allow for mechanistic research into human genetic variants implicated in advanced NAFLD.

## Results

### Human hepatocytes in chimeric mice on Western diet rapidly develop steatosis

To investigate how the human graft in liver chimeric mouse models responds to a hypercaloric diet, we first created humanized mice by transplanting primary human hepatocytes (PHH) into preconditioned immunocompromised *Fah^−/−^* NOD *Rag1^−/−^ Il2rg^null^* (FNRG) mice as described previously.^24,25^ Cycling the protective drug nitisinone^26^ resulted in at least 50% humanization after which mice were subjected to an *ad libitum* Western-style diet (WD), which consists of 60% fat in diet and 10% sucrose in drinking water (**Fig 1a**). WD feeding did not result in graft loss as determined by persistently high^27–29^ human serum albumin (hAlb) levels (**Suppl Fig S1a**). Periodic examination of huFNRG mouse livers showed steatosis^30^ as early as 4 weeks on WD as evidenced by hematoxylin and eosin (H&E) staining (**Fig 1b**) and Oil Red O staining (**Suppl Fig S1b**). Steatosis almost exclusively affected the human graft whereas the remaining mouse hepatocytes, which appear larger with darker cytoplasm after H&E staining, rarely developed steatosis. The contrast in steatosis between mouse and human hepatocytes was further evident by using a human-specific marker (**Fig 1c**). Livers from highly humanized mice with severe steatosis showed sparing around periportal regions (**Fig 1b**), reminiscent of zonal steatosis in adult NAFLD patients. The zonal predominance was further highlighted by glutamine synthetase staining, which is mostly expressed in zone 3 hepatocytes (**Fig 1d**). In order to quantify steatosis in huFNRG mice, the human areas of chimeric livers were scored blindly according to clinical criteria.^30^ All animals developed mild to severe steatosis as early as 4 weeks on WD. Steatosis persisted for 20 weeks on WD, which contrasted with huFNRG mice on control chow (chow) that never developed more than 5% steatosis (**Fig 1e**). Clinical scoring was confirmed by total triacylglycerol (TAG) quantification in huFNRG livers, which was 7 to 11-fold higher in WD than chow mice (**Fig 1f**), and liver total cholesterol (**Suppl Fig S1c**). Overall, the accumulation of TAG in huFNRG livers correlated to the steatosis score of the human graft (**Suppl Fig S1d**). These findings show that human hepatocytes in huFNRG mice rapidly developed moderate steatosis in response to WD feeding.

**Figure 1:**
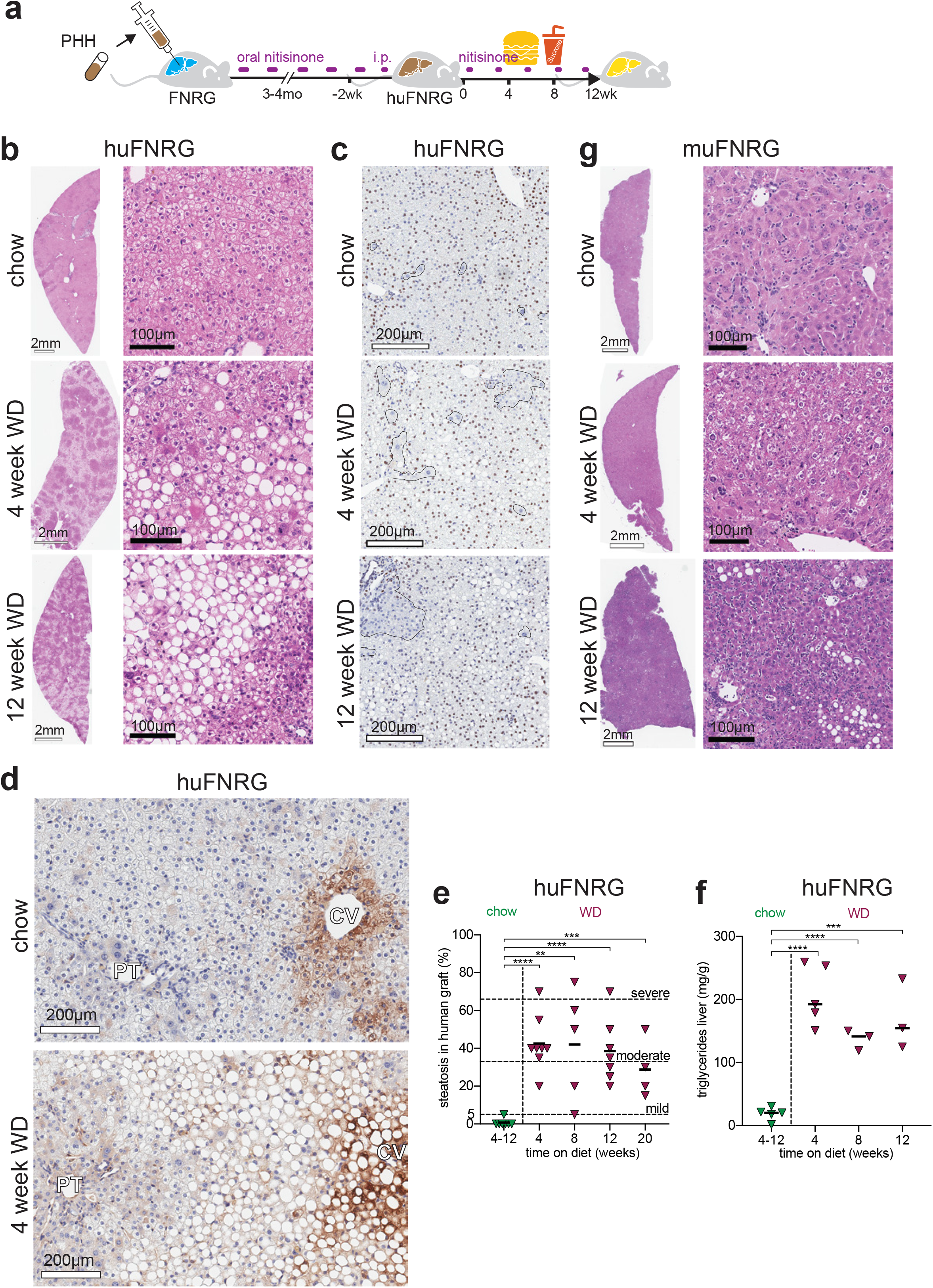
Human hepatocytes in chimeric mice on WD rapidly develop steatosis. **a)** Experimental timeline of FNRG mice that were transplanted with primary human hepatocytes (PHH), cycled off nitisinone to create huFNRG mice, and then subjected to chow or WD for up to 12 weeks. **b)** H&E staining on livers from huFNRG mice on chow and after 4 or 12 weeks on WD. **c)** Staining against human nuclear mitotic apparatus-1 in livers of huFNRG mice on chow and 4 or 12 weeks on WD. Areas with predominantly mouse hepatocytes are outlined. **d)** Staining against human glutamine synthetase on livers from huFNRG mice on chow and 4 weeks on WD. Central vein (CV), portal tract (PT). **e)** Liver steatosis score of the human graft in huFNRG mice on chow and after 4 to 20 weeks on WD. Symbols individual mice, bars are median, unpaired *t*-test with **p<0.005, ***p<0.0005 and ****p<0.0001. **f)** Hepatic triglyceride levels in livers of huFNRG mice on chow and after 4 to 12 weeks on WD. Symbols individual mice, bars are median, unpaired *t*-test with ***p<0.0005 and ****p<0.0001. **g)** H&E staining on livers from muFNRG mice on chow and after 4 or 12 weeks on WD.

We next tested if the marked steatosis difference between human and mouse hepatocytes was due to the immunodeficient mouse background, the intermittent nitisinone used to protect *Fah^−/−^* mice from liver injury,^31^ or the hepatocyte transplantation biology of the FNRG model. First, we subjected immunodeficient NOD *Rag1^−/−^ Il2rg^null^* (NRG) mice to WD feeding and nitisinone cycling. NRG livers developed minimal steatosis even after 12 weeks on WD (**Suppl Fig S1e**). To test if FNRG liver injury and/or hepatocyte repopulation physiology made hepatocytes susceptible to hypercaloric feeding we transplanted NRG mouse hepatocytes into FNRG mice. These animals were designated ‘murinized’ FNRG (muFNRG) mice. After 4 weeks on WD muFNRG livers did not display steatosis, but small areas of hepatocyte steatosis were present after 12 weeks of WD feeding (**Fig 1g**) which was in line with liver TAG levels (**Suppl Fig S1f**). These results show that human hepatocytes responded differently to WD feeding than mouse hepatocytes in these models.

We then tested if rapid steatosis was specific to this human hepatocyte donor (PHH1) and to the FNRG liver injury model, extending our protocols to other donors and liver chimeric mouse models. We first transplanted PHH2 into FNRG mice and thymidine kinase transgenic (TK-NOG) liver injury mice.^32^ Despite reaching lower levels of humanization in both models, human hepatocyte islands developed steatosis in response to WD feeding but not on chow (**Suppl Fig S1g**). We also transplanted a third PHH3 donor into urokinase plasminogen activator (uPA)/SCID mice^33^ and FNRG mice. Interestingly, uPA/SCID mice showed mild steatosis in the human graft on chow, which contrasted to the absence of steatosis in huFNRG livers on chow (**Suppl Fig S1h**). In the huFNRG model, PHH3 hepatocytes developed steatosis after 4 weeks of WD feeding. These data illustrate that humanized livers rapidly became steatotic following WD feeding regardless of the PHH donor, with different requirements for hypercaloric feeding between liver injury models.

Combined, these results demonstrate that human hepatocytes but not mouse hepatocytes in liver chimeric mice developed moderate steatosis as early as 4 weeks on WD.

### Western diet affects systemic metabolic parameters of chimeric FNRG mice

NAFLD clinically overlaps with obesity, dyslipidemia, insulin resistance and possibly hypertension.^2^ To determine whether WD affects metabolic parameters in liver chimeras, we first measured body weight accumulation in adult huFNRG mice on an *ad libitum* WD or control chow (chow) diet. Compared to chow, WD caused nearly 20% increase in body weight within 4 weeks on the diet and maintained this difference through 12 weeks on WD (**Fig 2a**). Furthermore, the gonadal fat fraction (**Fig 2b**) increased 2- to 2.4-fold in huFNRG mice on WD compared to chow. These data demonstrate that WD feeding resulted in weight gain and gonadal fat accumulation in huFNRG mice.

**Figure 2:**
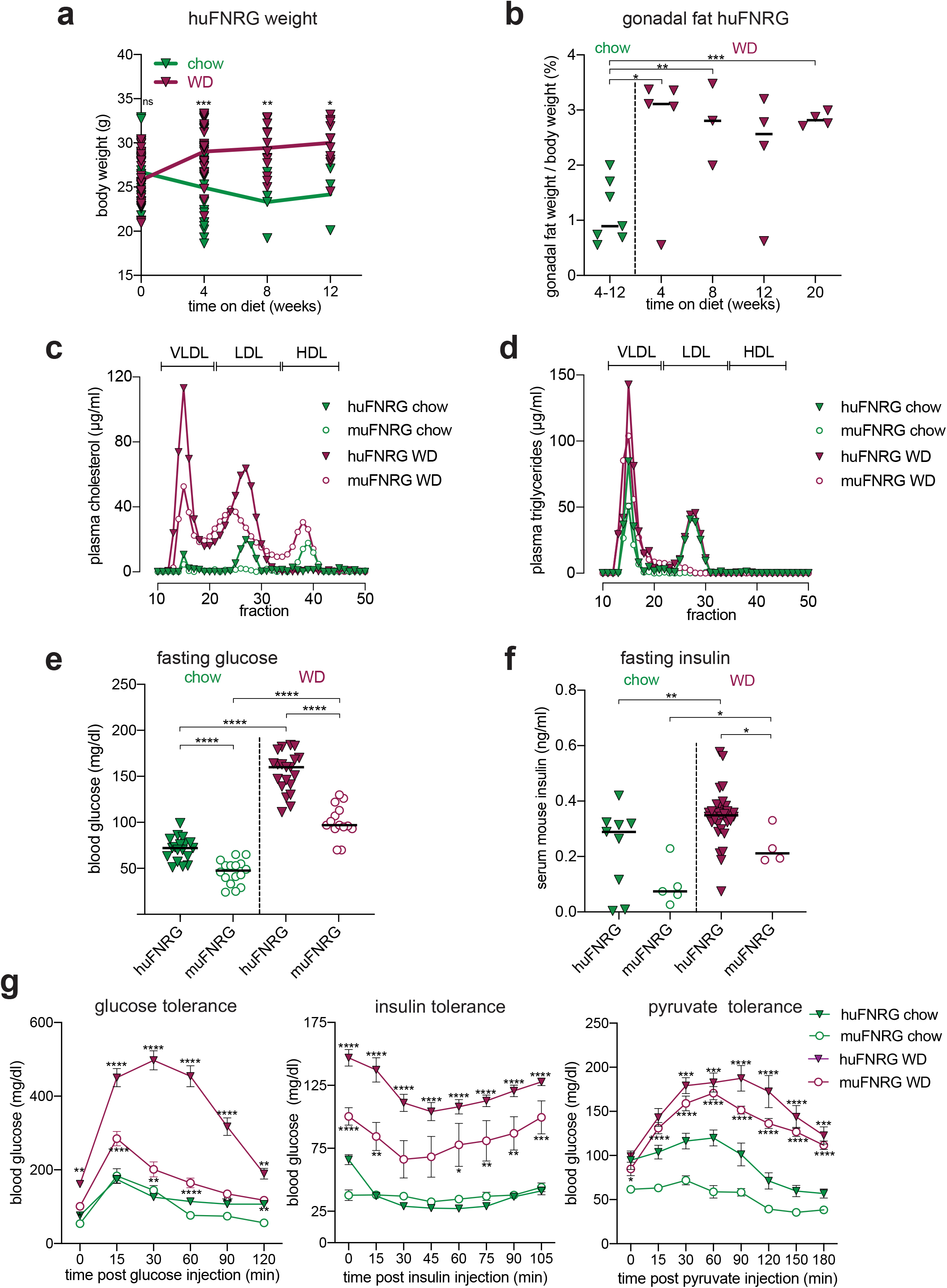
Systemic metabolic effects of WD in chimeric FNRG mice. **a)** Body weight in huFNRG mice during time course of chow or WD feeding. Symbols individual mice, line is mean, two-way ANOVA Sidak pairwise comparison *p<0.05, **p<0.005, ***p<0.0005. **b)** Gonadal fat fraction of total body weight in huFNRG mice on chow and after 4 to 20 weeks on WD. Symbols individual mice, bars are median, unpaired *t*-test with *p<0.05, **p<0.005, ***p<0.0005. **c)** Cholesterol in plasma lipoprotein fractions from huFNRG and muFNRG mice on chow and after 4 weeks on WD. Pooled plasma (3 mice/group), symbols are technical means. **d)** Triglycerides in plasma lipoprotein fractions from huFNRG and muFNRG mice on chow and after 4 weeks on WD. Pooled plasma (3 mice/group), symbols are technical means. **e)** Fasting blood glucose from huFNRG and muFNRG mice on chow and after 4 weeks on WD. Symbols individual mice, lines are median, unpaired *t*-test ****p<0.0001. **f)** Fasting serum mouse insulin from huFNRG and muFNRG mice on chow and after 4 weeks on WD. Symbols individual mice, lines are median, unpaired *t*-test with *p<0.05, **p<0.005. **g)** Intraperitoneal glucose, insulin and pyruvate tolerance testing in huFNRG and muFNRG mice on chow and after 4 weeks on WD. Blood glucose values measured at baseline (0) and at indicated timepoints after i.p. injections with each agent. Symbols are mean ± SEM of 5-7 mice per group, two-way ANOVA Sidak pairwise comparison of huFNRG on chow and WD (p-values on top) or muFNRG on chow and WD (p-values in bottom), *p<0.05, **p<0.005, ***p<0.0005 and ****p<0.0001

We then tested plasma lipids in chimeric mice. Hepatocytes are central to lipid homeostasis, and humanized mice have plasma cholesterol profiles shifted toward lower density lipoproteins compared to wild type mice.^34^ Similar to previous reports,^35–37^ huFNRG on chow contained more LDL than HDL cholesterol particles compared to muFNRG mice (**Fig 2c**). Four weeks of WD feeding resulted in stark increases in VLDL and LDL cholesterol in huFNRG plasma. Notably, the HDL fraction in huFNRG mice decreased with WD feeding whereas it rose in WD-fed muFNRG mice compared to chow (**Suppl Fig S2a**). Relative to chow, WD increased fasting serum triglycerides fractions, mostly in VLDL, and to a lesser extent in LDL fractions in huFNRG mice (**Fig 2d, Suppl Fig S2b**). These data show that WD feeding of huFNRG mice increased lower density lipoproteins and decreased HDL.

Given the strong clinical overlap between NAFLD and type 2 diabetes, we tested the effect of WD on glucose tolerance in huFNRG mice relative to muFNRG animals. While humanized animals started at a higher baseline, four weeks on WD resulted in a 2.2-fold increase in fasting blood glucose both in huFNRG and muFNRG mice (**Fig 2e)**. Fasting mouse insulin levels on chow appeared higher in humanized than murinized FNRG animals, and increased only slightly in huFNRG animals after 4 weeks on WD (**Fig 2f**). To further characterize glucose metabolism, fasting animals were subjected to glucose, insulin or pyruvate tolerance tests. After 4 weeks on WD, huFNRG mice displayed severely impaired glucose clearance with 3.1-fold higher cumulative blood glucose than chow, whereas this difference was 1.7-fold higher in muFNRG mice (**Fig 2g**). By contrast huFNRG mice on WD remained responsive to insulin challenge (**Fig 2g**). To assess gluconeogenesis in these animals, a pyruvate injection after 16 hours fasting resulted in a sharper rise in blood glucose in huFNRG on a WD than chow (**Fig 2g**). Eight weeks of WD feeding further prolonged time for glucose clearance in huFNRG mice (**Suppl Fig S2c**) without affecting endogenous insulin secretion (**Suppl Fig S2d**). These data show that WD feeding of chimeric FNRG mice affected glucose homeostasis without altering insulin-mediated glucose uptake.

Together these data show that in huFNRG mice WD causes increased body weight, dyslipidemia and impaired glucose metabolism compared to chow-fed animals.

### Western diet causes mild steatohepatitis in huFNRG mice

Non-alcoholic steatohepatitis (NASH) frequently causes elevated serum transaminases^38^ and is diagnosed histologically by the presence of hepatocyte ballooning and mononuclear infiltrates in the setting of >5% hepatocyte steatosis.^30^ Because the human graft rapidly developed steatosis, we tested whether huFNRG mice on WD also developed features associated with NASH.

We quantified the activities of serum transaminases ALT and AST, which were elevated in huFNRG mice 4 weeks on WD compared to chow. With prolonged WD feeding, ALT activity remained high while AST increased further (**Fig 3a**). To determine whether the ALT was produced by human or mouse hepatocytes, human-specific ALT protein was quantified and this correlated with serum ALT activity (**Suppl Fig S3a**). The human origin of transaminase activity was further supported by the lack of ALT activity and only minimal increases in AST activity in muFNRG mice on WD (**Suppl Fig S3b**). These data show that WD increased human transaminases in huFNRG serum.

**Figure 3:**
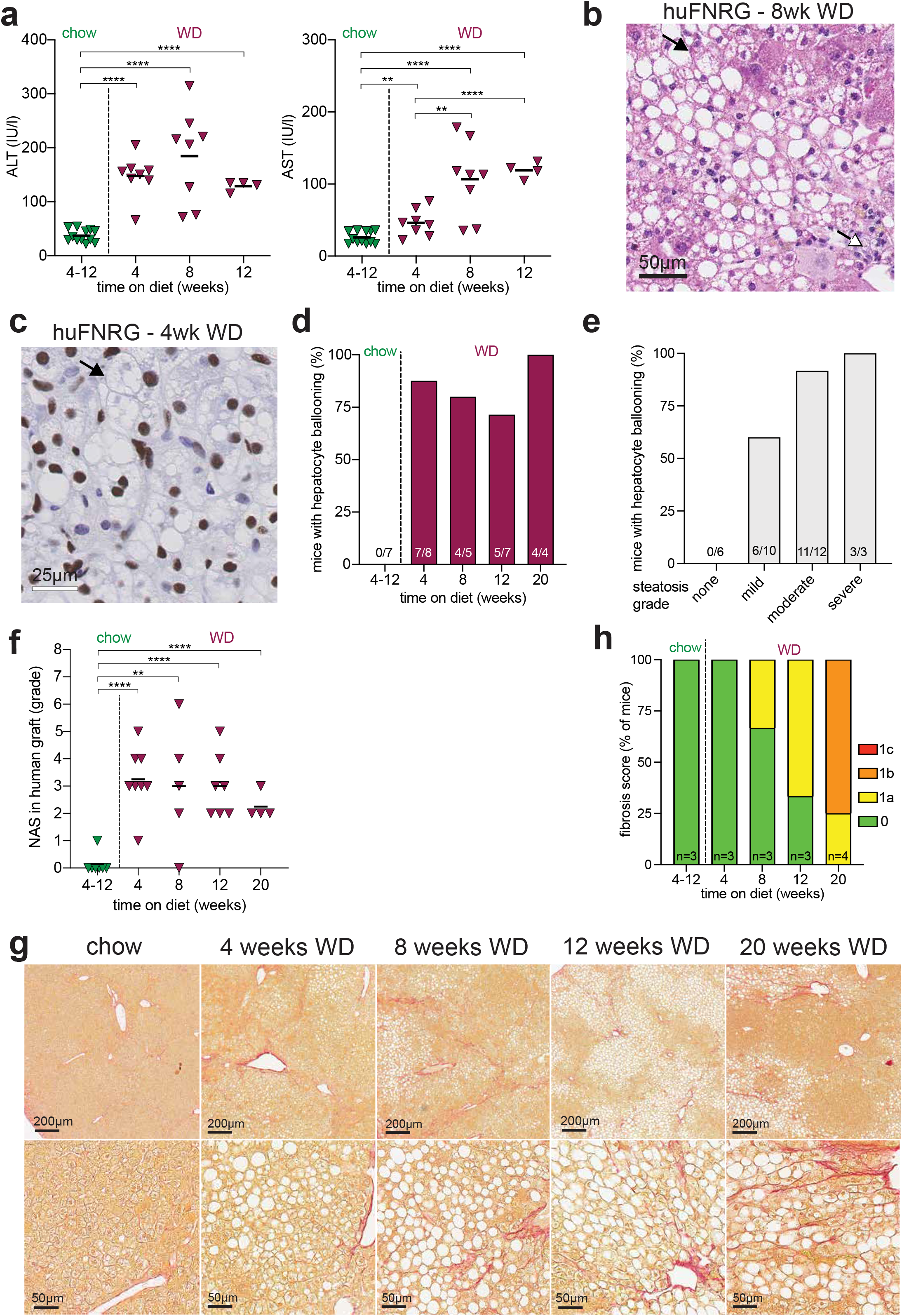
huFNRG mice on WD develop mild steatohepatitis. **a)** ALT and AST activity in serum from huFNRG mice on chow and after 4 to 12 weeks on WD. Symbols individual mice, bars are median, unpaired *t*-test **p<0.005, ****p<0.0001. **b)** H&E staining from a huFNRG mouse liver after 8 weeks on WD. Black arrow indicates hepatocyte ballooning degeneration and white arrow indicates lobular inflammation. **c)** Staining against human nuclear mitotic apparatus-1 in the liver of a huFNRG mouse after 4 weeks on WD. Black arrow indicates hepatocyte ballooning. **d)** Fractions of huFNRG mice with human hepatocyte ballooning degeneration on chow and WD over time. Number of mice with ballooning per group at bottom of bars. **e)** Fractions of huFNRG mice with human hepatocyte ballooning degeneration by grade of steatosis. Number of mice with ballooning per group at bottom of bars. **f)** NAFLD activity score (NAS) in the human graft of huFNRG mice on chow and after 4 to 20 weeks on WD. Symbols individual mice, bars are median, unpaired *t*-test **p<0.005, ****p<0.0001. **g)** Picrosirius Red staining for collagen in livers from huFNRG mice on chow and after 4 to 20 weeks on WD. **h)** Fibrosis stages in the human graft of huFNRG liver on chow and after 4 to 20 weeks on WD. Mouse numbers at bottom of bars.

We next assessed whether human hepatocytes in huFNRG mice on a WD displayed ballooning degeneration, a key characteristic of NASH.^30^ As early as 4 weeks on a WD, steatotic hepatocytes in huFNRG mice showed ballooning degeneration that persisted for 20 weeks (**Fig 3b, Suppl Fig S3c**). Hepatocyte ballooning was specific to the human graft (**Fig 3c**) and not donor hepatocyte-specific (**Suppl Fig S3d**). The fraction of huFNRG mice with ballooning degeneration did not increase with prolonged WD feeding (**Fig 3d**) but rather associated with the severity of steatosis grade (**Fig 3e**). Ballooning degeneration, which remains less well-defined in mouse hepatocytes,^19^ was not observed in either NRG or muFNRG mice on WD. Another key feature of NASH is the presence of lobular inflammation. After 4 weeks on WD, inflammatory infiltrates were detected in huFNRG mouse livers (**Fig 3b, Suppl Fig S3c**). This lobular inflammation was observed in approximately a third of huFNRG mice up to 12 weeks on WD but was not observed at 20 weeks (**Suppl Fig S3e**). We then scored the livers of huFNRG mice on WD according to the NAFLD activity score (NAS)^39^ within the human graft. After 4 weeks on WD, huFNRG mice developed borderline NASH that did not increase for up to 20 weeks on WD (**Fig 3f**). These data show that huFNRG mice developed mild NASH following WD feeding.

Pericellular fibrosis can accompany NASH and is thought to result from ongoing steatohepatitis.^40^ To test if WD feeding of huFNRG mice caused fibrosis, livers were stained for collagen with Picrosirius Red (PSR). Pericellular collagen depositions were first observed in a few mice after 8 weeks on WD (**Fig 3g**), then in a larger fraction at 12 weeks and finally more prevalent after 20 weeks on WD (**Fig 3h**). Trichrome staining confirmed collagen depositions around human hepatocytes, which are smaller with paler cytoplasm compared to mouse hepatocytes (**Suppl Fig S3f**). Fibrosis was not PHH1 specific since mice humanized with donor PHH2 showed a similar phenotype after 10 weeks on WD (**Suppl Fig S3g**). These data indicate that prolonged WD feeding results in mild fibrosis around human hepatocytes.

Overall, these findings show that huFNRG mice developed mild NASH as early as 4 weeks on WD, which was accompanied by pericellular fibrosis in the human graft after 8 weeks of WD feeding.

### WD affects human and mouse liver transcriptomes differentially over time

In huFNRG chimeras that were created with PHH the non-parenchymal cells and leukocytes remained murine (^27^ and unpublished observations). To determine how WD diet affected human and mouse gene expression we performed RNA-sequencing (RNA-seq) on livers from fasting huFNRG mice. STAR Alignment mapped >85% of transcripts uniquely to a human reference genome (**Suppl Fig S4a**).

Transcripts aligned to human genome (human reads) in mice that were fed WD for 4 weeks were compared to animals on chow, which identified 662 differentially expressed transcripts (**Fig 4a**). Among mice on WD human transcripts changed progressively less over time, totaling 154 at 8 vs. 4 weeks, 127 at 12 vs. 8 weeks, and 95 at 20 vs. 12 weeks (**Fig 4a**).

**Figure 4:**
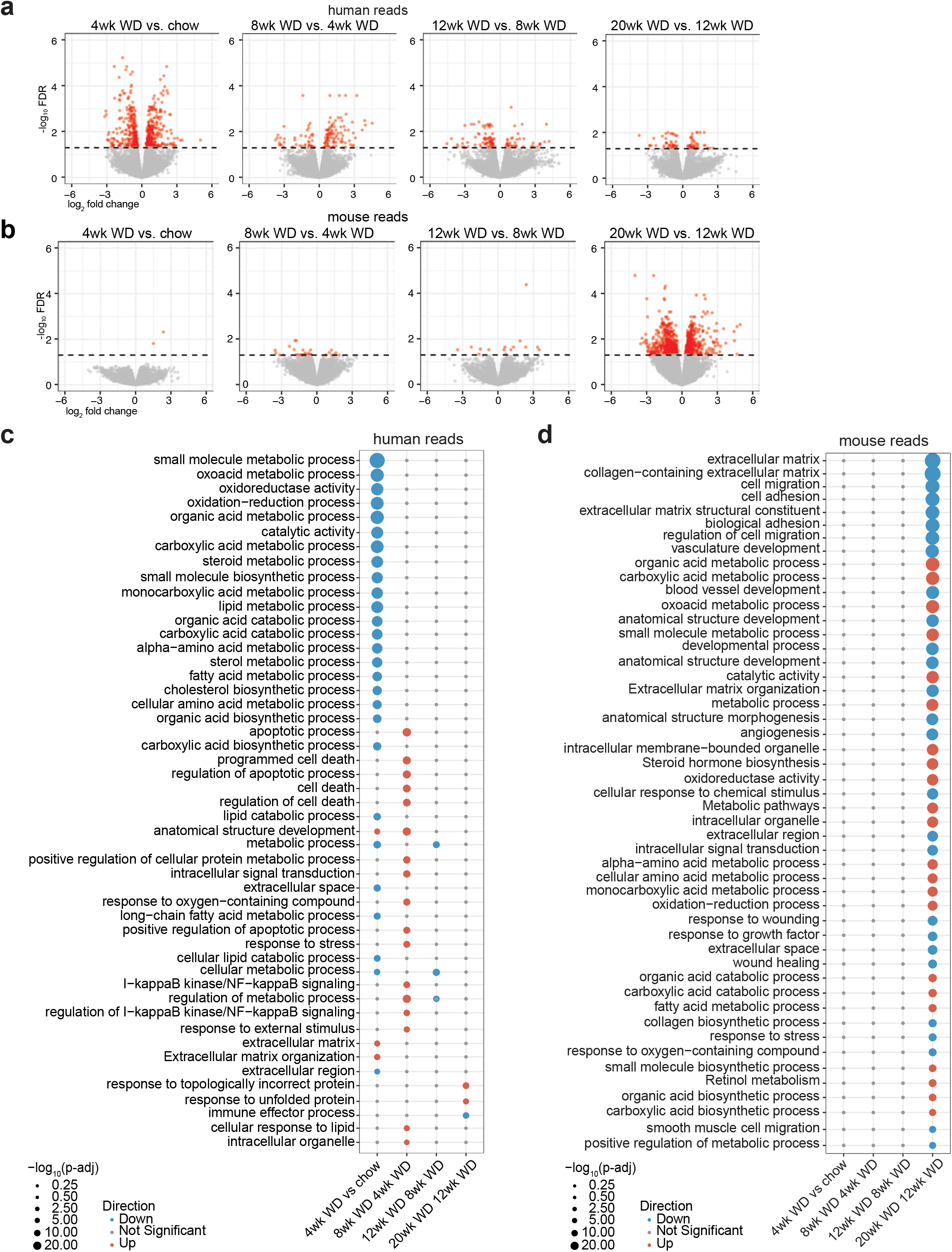
Transcriptional changes in huFNRG mice on WD. **a)** Volcano plots of liver transcripts mapped to human genome (human reads) comparing huFNRG mice 4 weeks on WD to chow, and longer WD durations to the previous WD timepoint. RNA-sequencing of 3-4 mice per group, red symbols FDR <0.05, grey denotes not significant. **b)** Volcano plots of liver transcripts mapped to mouse genome (mouse reads) comparing huFNRG mice 4 weeks on WD to chow, and longer WD durations to the previous WD timepoint. RNA-sequencing of 3-4 mice per group, red symbols FDR <0.05, grey denotes not significant. **c)** Gene-ontology (GO) pathway analysis of human reads from livers of huFNRG mice 4 weeks on WD to chow, and longer WD durations to the previous WD timepoint. Blue symbols are down- and red symbols are upregulated pathways, grey denotes no significant changes, symbol size indicates statistical significance. **d)** Gene-ontology (GO) pathway analysis of mouse reads from livers of huFNRG mice 4 weeks on WD to chow, and longer WD durations to the previous WD timepoint. Blue symbols are down- and red symbols are upregulated pathways, grey denotes no significant changes, symbol size indicates statistical significance.

We then aligned transcripts of these highly humanized livers to mouse reference genome (mouse reads). In contrast, only 2 mouse transcripts were statistically different between huFNRG mice that were fed WD or chow for 4 weeks. Transcriptional changes in the mouse reads remained low for the first 12 weeks of WD feeding, but a large number of differentially expressed transcripts were observed by 20 weeks (**Fig 4b**).

To further investigate changes in the gene expression profile we performed gene ontology (GO) analysis using the GO, Kyoto Encyclopedia of Genes and Genomes (KEGG), REACTOME (REAC), and Transcription Factor (TF) databases. GO (**Fig 4c)**, KEGG and REAC (**Suppl Fig S4b**) alignments of human reads revealed mostly downregulation of metabolic and catabolic pathways in mice 4 weeks on WD compared to chow, while transcription factors were upregulated (**Suppl Fig S4b**). A notable exception to many downregulated GO pathways was the Extracellular Matrix pathway, which was upregulated at 4 weeks of WD feeding. Comparing 8 vs. 4 weeks on WD identified upregulation of human GO pathways involved in cell death and NF-κB signaling. Later timepoints revealed fewer and statistically less significant pathways. Few mouse GO pathways were changed by WD feeding at early time points. At 20 weeks many pathways involved in cellular development and metabolism changed (**Fig 4d, Suppl Fig S4c**). Notably several metabolic pathways that were downregulated in the human transcriptome at 4 weeks were upregulated in the mouse transcriptome at 20 weeks.

Taken together, these data indicate that many metabolic pathways were transcriptionally downregulated in the human graft as early as 4 weeks on WD, while the mouse transcriptome of huFNRG mice remained largely silent until 20 weeks of WD feeding.

### Mice humanized with a homozygous PNPLA3-148M donor develop steatohepatitis

Thus far huFNRG mice were humanized with PHH homozygous for PNPLA3-148I (148I-huFNRG). We next set out to test how chimeric mice humanized with the PNPLA3-148M variant responded to WD. To this end *Fah^−/−^ Rag2^−/−^ Il2rg^null^* (FRG) mice^26^ were humanized with a homozygous PNPLA3-148M donor (148M-huFRG) and given WD for 4 weeks. As expected WD caused metabolic changes, including raised fasting blood glucose, a minimal increase in fasting insulin and generally higher plasma triglycerides (**Suppl Fig S5a-c**). Livers of 148M-huFRG mice on WD had moderate (>33%) and severe (>66%) steatosis in contrast to <5% steatosis that was observed in chow-fed mice (**Fig 5a-b**). Histological findings were confirmed by 5.3-fold increase in TAG concentrations in the livers of 148M-huFRG mice on WD compared to chow-fed controls (**Suppl Fig S5d**). In contrast to previous 148I-huFNRG studies, the human graft showed not only macrovesicular but also microvesicular steatosis in all 148M-huFRG mice on WD (**Fig 5c**). In addition, 148M-huFRG livers contained Mallory-Denk bodies, high grade of ballooning degeneration and lobular inflammation, resulting in a median NAS of 5.2 (**Fig 5d**). These data demonstrate that 148M-huFRG mice subjected to WD for 4 weeks showed features of NASH.

**Figure 5).**
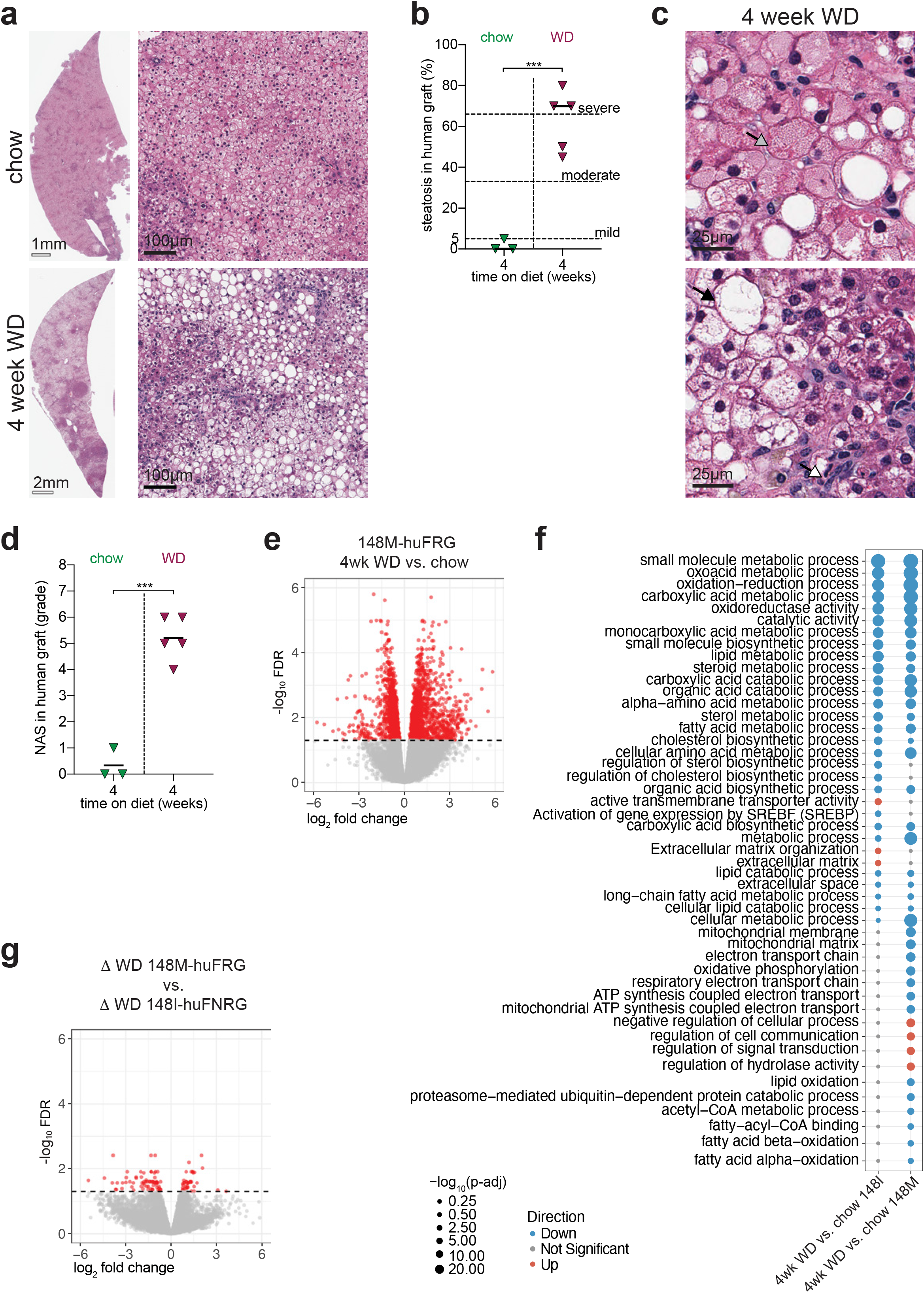
PNPLA3-148M huFRG mice on WD develop steatohepatitis. **a)** H&E staining on livers from 148M-huFRG mice on chow and after 4 weeks on WD. **b)** Clinical steatosis score of the human graft on livers from 148M-huFNRG mice on chow and after 4 weeks WD. Symbols are individual mice, bars median, unpaired *t*-test ***p<0.0005. **c)** H&E staining on livers from 148M-huFRG mice 4 weeks on WD. Grey arrow indicates Mallory Denk body, black arrow ballooning degeneration and white arrow lobular inflammation. **d)** NAFLD Activity Score (NAS) in the human graft of 148M-huFRG mice on chow and after 4 weeks WD. Symbols individual mice, bars are median, unpaired *t*-test ***p<0.0005. **e)** Volcano plots of liver transcripts mapped to human genome (human reads) comparing 148M-huFRG mice after 4 weeks WD vs. chow. RNA-sequencing of 3-4 mice per group, red symbols FDR <0.05, grey denotes not significant. **f)** Gene-ontology (GO) pathway analysis of human reads comparing 148I-huFNRG and 148M-huFRG livers after 4 weeks WD vs. chow. Blue symbols are down- and red symbols are upregulated pathways, grey denotes no significant changes, symbol size indicates statistical significance. **g)** Volcano plots of human reads differentially expressed after 4 weeks WD compared to chow (∆ WD) in 148M-huFRG livers versus 148I-huFNRG livers. Red symbols FDR <0.05, grey denotes not significant.

To investigate if 148M-huFRG mice transcriptionally responded differently to WD feeding than 148I-huFNRG animals, RNA-seq analyses was performed. As expected, the human graft in 148M-huFRG mice responded to 4 weeks on WD with numerous transcriptomic changes compared to chow, with 1458 downregulated and 1294 upregulated genes (**Fig 5e**). Pathway analyses revealed that metabolic and catabolic pathways were mostly downregulated by WD in 148M-huFRG mice, similar to previous results in 148I-huFNRG mice. Despite the similarities between both models, there were pathways, including oxidative phosphorylation and mitochondrial function, which were downregulated following WD only in the livers of 148M-huFRG mice but not in 148I-huFNRG. In addition, some pathways were differentially upregulated by WD between these two models (**Fig 5f**). Comparing individual genes from livers of 148M-huFRG mice and 148I-huFNRG, there were 2329 differently expressed transcripts on chow and 3268 on WD (**Suppl Fig S5e**). However, when comparing transcripts that changed by WD feeding over chow in each model (Δ WD), there were only 108 genes differentially regulated between these models (**Fig 5g**).

Taken together, these findings show that 148M-huFRG mice developed microvesicular steatosis and NASH after 4 weeks on WD, while few pathways were transcriptionally different from 148I-huFNRG mice on the same diet.

### PNPLA3-148M overexpression in hepatocytes exacerbates steatosis

In addition to the PNPLA3 genotype, 148M-huFRG and 148I-huFNRG mice were distinct in numerous other human genetic and mouse background variables. To better characterize if phenotypic differences were due to the PNPLA3 variant, we transduced mouse-passaged PHH cultures from a 148I homozygous donor (PHH1) with PNPLA3-148M and red fluorescent protein (RFP), 148I and RFP or RFP only before re-transplantation into mice.^25^ Humanization kinetics did not differ between the PNPLA3 variants or RFP controls as determined by serum hAlb values (**Suppl Fig S6a**). Once hAlb serum levels plateaued, huFNRG mice with transduced PNPLA3-148M hepatocytes (td148M), td148I or tdRFP mice were subjected to 4 weeks on WD or chow (**Fig 6a**).

**Figure 6:**
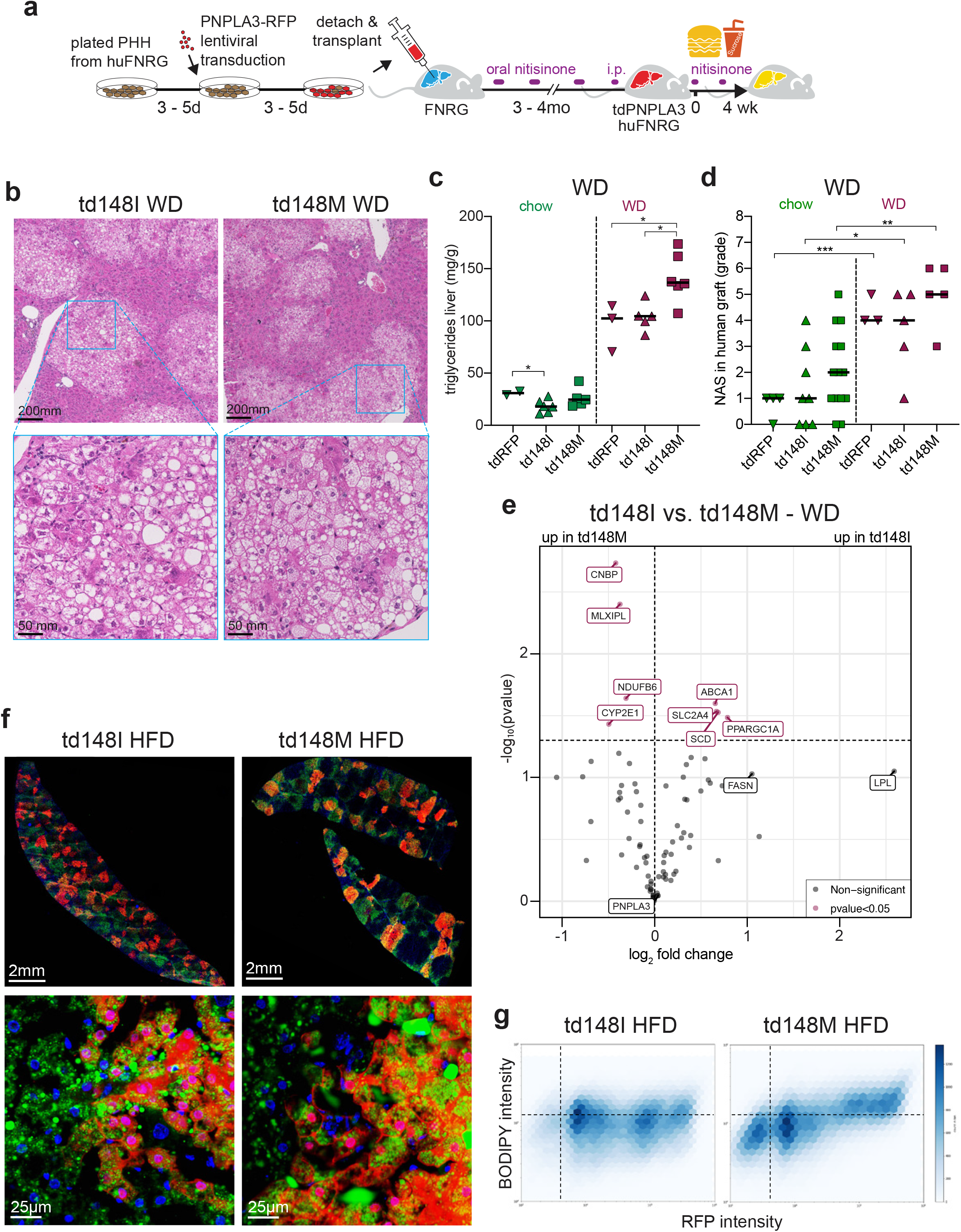
Hepatocyte PNPLA3-148M overexpression worsens steatosis. **a)** Experimental timeline of creating PNPLA3 overexpressing mice and diet challenge. Mouse passaged primary human hepatocytes (PHH) cultures were transduced with lentiviral vectors expressing PNPLA3 variants and re-transplanted. Following humanization, huFNRG with PNPLA3 transduced hepatocytes (td) were exposed to 4 weeks of hypercaloric feeding. **b)** H&E staining on livers from td148I and td148M mice after 4 weeks on WD. **c)** Hepatic triglyceride quantification in tdRFP, td148I and td148M mice on chow and after 4 weeks on WD. Symbols individual mice, bars are median, unpaired *t*-test, *p<0.05. **d)** NAFLD Activity Score (NAS) in the human graft of tdRFP, td148I and td148M mice on chow and after 4 weeks on WD. Symbols individual mice, bars are median, unpaired *t*-test, *p<0.05, **p<0.005, ***p<0.0005. **e)** Volcano plot of 80 human genes expressed in livers of td148I versus td148M mice after 4 weeks on WD. Expression by qRT-PCR normalized to 11 housekeeping genes, n=3-4 mice per group, burgundy symbols p<0.05, grey symbols not statistically significant. **f)** Low and high magnification fluorescent images of livers from td148I and td148M mice after 4 weeks of HFD stained for neutral lipids (green, BODIPY) and nuclei (blue, DAPI). **g)** Two-dimensional density plots of neutral lipids (BODIPY) and RFP of 355,161 small hepatocytes in livers from four td148I mice and 183,289 small hepatocytes in livers from three td148M mice after 4 weeks of HFD.

WD induced human hepatocyte steatosis in all three groups of mice, with only td148M livers containing widespread microvesicular steatosis (**Fig 6b, Suppl Fig S6b**). Strikingly, even on chow td148M livers contained rare areas with human microvesicular steatosis, which was not observed in td148I or tdRFP livers (**Suppl Fig S6c)**. Lipid quantification in livers from mice on WD showed 36% more triglycerides in td148M livers compared to td148I or tdRFP (**Fig 6c**). Despite these phenotypic changes, the NAS was not statistically different in td148M mice from td148I or tdRFP animals (**Fig 6d**). These data demonstrate that PNPLA3 overexpressing huFNRG mice on WD developed NASH, with td148M livers containing widespread microvesicular steatosis.

Since RNA-seq pathway analyses did not identify many differences between the 148M-huFRG and 148I-huFNRG models (**Fig 5g**), we performed qRT-PCR for 80 human genes associated with fatty liver diseases. Liver tissue from td148I and td148M mice that were fed chow revealed 20 genes that were expressed differently, with lipoprotein lipase showing the greatest difference (**Suppl Fig S6d**). Comparing both groups following WD we identified only 8 genes that were affected by 2-fold or less, of which *CYP2E1* was most transcriptionally upregulated in td148M livers (**Fig 6e**). These results illustrate the modest expression differences between mice overexpressing the two PNPLA3 variants on WD.

We hypothesized that the minimal differences between td148M and td148I mice could be explained by the severe phenotype caused by WD, similar to what was reported in human PNPLA3 transgenic mice.^10^ Therefore, we challenged animals with the high fat diet (HFD, 60% fat) only, excluding the sucrose drinking water portion of WD. Four weeks of HFD resulted in mild triglyceride accumulation in the livers of td148I mice, which was exacerbated 3.2-fold in the livers of td148M mice (**Suppl Fig S6e**). In td148M mice the HFD resulted in widespread microvesicular steatosis and Mallory Denk bodies (**Suppl Fig S6f**). Low magnification fluorescence imaging of liver cross-sections stained for neutral lipids suggested that td148M livers on HFD contained more lipids than td148I mice (**Fig 6f**). At higher magnification steatosis appeared more severe in RFP positive areas of td148M livers whereas lipid distribution in td148I mice was not associated with RFP. We then analyzed RFP and neutral lipid intensity from individual small hepatocytes, calibrated to a highly humanized huFNRG mouse whose graft was severely steatotic (**Suppl Fig S6g**). A two-dimensional density plot of individual hepatocytes from three td148M animals showed that hepatocytes with higher RFP intensity contained more lipids, whereas lipids were equally distributed along RFP intensity in hepatocytes from four td148I mice (**Fig 6g**). These findings suggest that 148M but not 148I overexpression, as determined by an RFP reporter, exacerbated steatosis. These combined data show that PNPLA3-148M overexpression in hepatocytes from a 148I donor caused microvesicular steatosis and exacerbated steatosis.

## Discussion

In this study, we have established systems to model NAFLD genetics in human hepatocytes. The most striking phenotype in the ‘standard’ 148I-huFNRG model was the rapid onset of human steatosis on WD. This contrasted to the minimal steatosis in the residual mouse hepatocytes of huFNRG mice or in muFNRG controls. Both resembled non-humanized mouse models that require prolonged WD feeding to develop moderate steatosis.^41^ Impaired cross-talk between human hepatocytes and murine cells likely contributed, as we previously showed for FGF15 and FGF19 signaling,^42^ yet mouse and human hepatocytes may also inherently respond differently to overnutrition. This is supported by the clinical observations of how rapidly hepatic fat content changes depending on the type and amount of food intake. For example, a three-week overfeeding study increased intrahepatic triglycerides up to 55%^43^ and a carbohydrate-restricted diet reduced hepatic fat by 44% in two weeks.^44^ These clinical data illustrate that hepatic steatosis dynamically responds to nutritional variations. huFNRG mice can model these observations in humans and provide a platform to explore how dietary components affect human hepatocyte lipid compositions.

Hepatocytes are central to systemic lipid and glucose regulation. Similar to previous reports^35–37^ huFNRG on chow had starkly different lipid profiles than muFNRG, in part due to mice lacking cholesterol ester transfer protein.^37^ In addition, we show that fasting glucose and possibly fasting insulin levels were higher in huFNRG mice than muFNRG mice on chow. Species incompatibilities may contribute to the elevated fasting glucose of huFNRG mice on chow, as was illustrated by studies in the uPA liver chimeric model.^45^ While some human donors develop spontaneous steatosis in these chimeras, similar to what we observed with PHH3, administration of human growth hormone reversed this phenotype, possibly through IGF-1 signaling in the liver.^45,46^ We speculate that hepatocytes in huFNRG are likewise predisposed to steatosis, and therefore may only require a short duration of hypercaloric feeding to develop moderate steatosis. Interestingly, muFNRG controls without hepatic steatosis developed glucose intolerance on WD that was proportionally similar to huFNRG mice with moderate steatosis. Uncoupling hepatocyte steatosis from glucose homeostasis in these models may advance insights into clinically distinct forms of NAFLD with and without insulin resistance.^47^

Despite their broad immunodeficiency huFNRG mice displayed all the features that define NASH, including ballooning degeneration and lobular infiltrates. This illustrates that mild steatohepatitis and pericellular fibrosis can be initiated without lymphocytes. The cellular immune contributions to NASH pathogenesis remain poorly characterized. In human NAFLD, portal immune infiltrates contain macrophages and T cells whereas lobular infiltrates are dominated by macrophages and neutrophils.^48^ How these cell types and their subsets affect human NASH remains undefined and proposed mechanisms are based on animal models with unclear translational value.^49–51^ Some compounds that effectively interfered with the immunopathogenesis of murine steatohepatitis failed in NASH patients,^52^ and no immunological interventions have yet shown clear benefits in later phase clinical trials. Furthermore, species differences preclude investigations into non-conserved pathways, e.g. IL-32 that lacks a mouse orthologue. This cytokine was recently associated with advanced NAFLD^53^ and *IL32* was transcriptionally upregulated in 148I-huFNRG livers after 12 weeks and 148M-huFRG livers after 4 weeks of WD feeding. Going forward, human immune system mice can be combined with liver chimeras^27,54^ for studies into immune pathways that differ between mice and humans.

Transcriptomic characterization of huFNRG livers showed a large number of human metabolic pathways that were transcriptionally downregulated after 4 weeks of WD feeding and only a few pathways that were upregulated. Eight weeks of WD feeding caused upregulation of a small number of cellular stress and cell death pathways, and remarkably few human pathways transcriptionally changed after that. Equally surprising were the small number of mouse transcripts that changed during the first 12 weeks on WD. This may be partly explained by under-sampling of mouse hepatocytes at earlier timepoints, as illustrated by changes in mouse metabolic pathways after 20 weeks WD when humanization histologically appeared to decline. The murine non-parenchymal transcriptome, however, was also largely unperturbed by WD feeding. Whether changes from WD were masked by cyclical nitisinone withdrawal or if non-parenchymal cells did not transcriptionally respond to signals from the human graft will require further studies, including in other chimera models without cyclical liver injury.

As we have shown, these systems will now allow for mechanistic studies into genetic variants that drive NAFLD. Many polymorphisms associated with elevated transaminases^55^ and most variants implicated in advanced NAFLD^56^ are in genes expressed in hepatocytes. As a proof-of-principle, we modeled the PNPLA3-148M polymorphism.^6^ Its prevalent allele frequency allowed for the identification of a high-quality homozygous 148M donor to create human chimeras. PNPLA3 overexpression then controlled for the plethora of additional genetic differences between PHH donors. By forced overexpression of 148M in homozygous 148I hepatocytes, we observed the development of microvesicular steatosis. This strongly suggests that this phenotype in 148M-huFRG mice is related to their PNPLA3 genotype. Differences between 148I and 148M had previously been identified in mouse hepatocytes and human cell lines,^10,11,13,15,16,57^ none of which displayed microvesicular steatosis. Whether this phenotype, which clinically correlates with advanced NASH,^58^ is specific to human hepatocytes will require further investigation. Species discordant findings are further supported by 148M-huFRG mice that developed NASH on a normal-cholesterol WD. High sugar diets exacerbated steatosis in *Pnpla3*-148M knock-in mice^11^ but only a high-cholesterol diet caused NASH.^12^ Such differences are not unique to PNPLA3. A loss-of-function polymorphism in *HSD17B13* was associated with less severe fatty liver disease^59^ yet *Hsd17b13^−/−^* mice were not protected from obesogenic or alcoholic liver injury.^60^ These observations illustrate the need for human models to unravel mechanisms that drive advanced NAFLD.

Taken together these results show the value of liver chimeras for studies of pathways that have been genetically implicated in human fatty liver diseases. Notwithstanding their broad implications to advance human hepatocyte research, several technical challenges remain. Identifying high-quality PHH donors with rare allele variants, e.g. in *TM6SF2* or *MBOAT7*,^61,62^ will remain difficult. Additionally, creating isogenic controls will require improvements in gene editing efficiency of primary human materials, either to disrupt genes or prime edit polymorphisms.^63^ An alternative strategy is to prime edit renewable cells such as pluripotent stem cells. However, hepatocyte-like cells derived from these sources have thus far failed to reliably and reproducibly humanize mouse chimera models.^64^ Overcoming these technical challenges would further advance the models presented here and generate ever more relevant systems to dissect the pathophysiology of human fatty liver diseases.

## Figure Legends

**Suppl Figure S1: Human hepatocyte steatosis in chimeric models**

**a)** Serum human albumin (hAlb) levels in huFNRG mice during the course of chow and WD feeding.

**b)** Oil Red O staining on livers from huFNRG mice on chow and after 4 weeks on WD.

**c)** Hepatic cholesterol quantification from huFNRG mice on chow and after 4 to 12 weeks on WD. Symbols individual mice, bars are median, unpaired *t*-test with *p<0.05, **p<0.005.

**d)** Hepatic triglyceride levels versus clinical steatosis scoring of the human graft in huFNRG mice on chow and after 4 to 12 weeks on WD. Symbols individual mice, line is simple linear regression, r^2^ Pearson correlation coefficient.

**e)** H&E staining on livers from NRG livers on chow and after 12 weeks on WD.

**f)** Hepatic triglyceride quantification from muFNRG mice on chow, and after 4 and 12 weeks on WD. Symbols individual mice, bars are median.

**g)** Liver sections from PHH2 huFNRG mice on chow and after 10 weeks on WD stained for human nuclear mitotic apparatus-1 (left). Liver sections from PHH2 huTK-NOG mice on chow and after 10 weeks on WD stained for H&E (right).

**h)** H&E staining on livers from PHH3 huFNRG and PHH3 hu-uPA/SCID mice livers. High magnification images of a huFNRG mouse liver on chow (upper left); after 4 weeks on WD (lower left); low and high magnification images of a hu-uPA/SCID mouse liver on chow (right).

**Suppl Figure S2: Systemic metabolic effects of WD feeding**

**a)** Cholesterol quantification in plasma lipoprotein fractions from huFNRG and muFNRG mice on chow and after 4 weeks on WD. Pooled plasma (3 mice/group), column bars are technical means ± SEM, unpaired *t*-test ****p<0.0001.

**b)** Triglyceride quantification in plasma lipoprotein fractions from huFNRG and muFNRG mice on chow and after 4 weeks on WD. Pooled plasma (3 mice/group), column bars are technical means ± SEM, unpaired *t*-test *p<0.05, ***p<0.0005) and ****p<0.0001.

**c)** Intraperitoneal glucose tolerance testing in huFNRG on chow and after 8 weeks on WD. Symbols are mean ± SEM of 5-8 mice per group, two-way ANOVA Sidak pairwise comparison with *p<0.05, **p<0.005, ***p<0.0005 and ****p<0.0001

**d)** Mouse insulin secretion during glucose tolerance testing in huFNRG mice on chow and after 8 weeks on WD. Symbols are mean ± SEM of 5-8 mice per group.

**Suppl Figure S3: Mild steatohepatitis in huFNRG mice on WD**

**a)** Human ALT protein versus ALT activity in serum of huFNRG mice on chow and WD. Symbols individual mice, line is simple linear regression, r^2^ Pearson correlation coefficient.

**b)** ALT and AST activity in serum of muFNRG mice on chow and after 4 to 12 weeks on WD. Symbols individual mice, bars are median, unpaired *t*-test with **p<0.005, ****p<0.0001.

**c)** H&E staining on livers from huFNRG mice after 4, 12 and 20 weeks on WD. Black arrow indicates hepatocyte ballooning degeneration and white arrow indicated lobular inflammation.

**d)** H&E staining on liver from a huFNRG mouse engrafted with PHH3 after 4 weeks on WD. Black arrow indicates hepatocyte ballooning degeneration.

**e)** Fractions of huFNRG mice with lobular inflammation on chow and after 4 to 20 weeks on WD. Number of mice with lobular inflammation per group at bottom of bars.

**f)** Masson trichrome staining for collagen on livers from huFNRG mice on chow and after 12 and 20 weeks on WD.

**g)** Picrosirius Red staining for collagen on a liver of huFNRG mouse engrafted with PHH2 after 10 weeks on WD.

**Suppl. Figure S4: Transcriptional changes in huFNRG mice on WD**

**a)** Spliced Transcripts Alignment to a Reference (STAR) Scores of transcripts from fasting huFNRG mice livers on chow and after 4 to 20 weeks on WD aligned to human reference genome. Numbers on left indicate individual mouse tags.

**b)** Manhattan plots comparing human pathways in livers from huFNRG mice 4 weeks on WD to chow, and longer WD durations to the previous WD timepoint. Gene-ontology (GO) with GO-MF (Molecular Function), GO-CC (Cellular Component), GO-BP (Biological Process), Kyoto Encyclopedia of Genes and Genomes (KEGG), REACTOME (REAC) and Transcription Factor (TF). Symbol size indicates percentage of statistically significant differently expressed transcripts aligned to a certain pathway, only pathways statistically significant enriched for a group of mice are displayed.

**c)** Manhattan plots comparing mouse pathways in livers from huFNRG mice 4 weeks on WD to chow, and longer WD durations to the previous WD timepoint. Gene-ontology (GO) with GO-MF (Molecular Function), GO-CC (Cellular Component), GO-BP (Biological Process), Kyoto Encyclopedia of Genes and Genomes (KEGG), REACTOME (REAC) and Transcription Factor (TF). Symbol size indicates percentage of statistically significant differently expressed transcripts aligned to a certain pathway, only pathways statistically significant enriched for a group of mice are displayed.

**Suppl Figure S5: 148M-huFRG mice develop steatohepatitis**

**a)** Fasting blood glucose from 148M-huFRG mice on chow and after 4 weeks on WD. Symbols individual mice, bars are median, unpaired t-test, **p<0.005.

**b)** Fasting serum mouse insulin in 148M-huFRG mice on chow and after 4 weeks on WD. Symbols individual mice, bars are median.

**c)** Fasting plasma triglycerides in 148M-huFRG mice on chow and after 4 weeks on WD. Symbols individual mice, bars are median.

**d)** Hepatic triglyceride quantification in 148M-huFRG mice on chow and after 4 weeks on WD. Symbols individual mice, lines are median, unpaired t-test, **p<0.005.

**e)** Volcano plots showing differently expressed transcripts mapped to human reference genome (human reads) from 148M-huFRG versus 148I-huFNRG livers on chow (left) and after 4 weeks on WD (right). Red symbols FDR <0.05, grey denotes not significant.

**Suppl Figure S6: PNPLA3 overexpressing huFNRG mice**

**a)** Serum human albumin (hAlb) levels in FNRG mice after transplantation with transduced PHH. Symbols are mean ± SEM of 3-16 mice.

**b)** H&E staining on liver from a tdRFP mouse after 4 weeks on WD.

**c)** H&E staining on livers from td148I and td148M mice on chow. White arrow indicates macrovesicular and black arrow microvesicular steatosis.

**d)** Volcano plot of 80 human genes expressed in livers of td148I versus td148M mice on chow. Expression by qRT-PCR, normalized to 11 housekeeping genes, n=3-4 mice per group, green symbols p<0.05, grey symbols not statistically significant.

**e)** Hepatic triglyceride quantification in livers from td148I and td148M mice after 4 weeks of HFD. Symbols individual mice, bars are median, unpaired *t*-test *p<0.05.

**f)** H&E staining on liver from a td148M mouse after 4 weeks of HFD. White arrow indicates lobular inflammation, black arrow microvesicular steatosis and grey arrow Mallory-Denk body.

**g)** H&E staining on liver of non-transduced huFNRG mouse with severe steatosis (left). Right panel shows two-dimensional density plot of neutral lipids (BODIPY) and RFP of small hepatocytes in liver from mouse displayed on left.

## Acknowledgements

This work was supported by the National Institutes of Health Grants R37DK048873, R01DK056626 and R01DK103046 (to DEC), R01DK085713 (to CMR), K08DK090576 and R01AA027327 (to YPJ) and the Starr Foundation (to CMR). PM was supported by grants from Ghent University (Special Research Fund – ICOH expert center) and the Research Foundation – Flanders (grants VirEOS 30981113 and G047417N). MK received funding from the Deutsche Forschungsgemeinschaft under grant number KA4688/1-1. The project was co-sponsored by the Center for Basic and Translational Research on Disorders of the Digestive System through the generosity of the Leona M. and Harry B. Helmsley Charitable trust (to EM). The NYULH Center for Biospecimen Research and Development, Histology and Immunohistochemistry Laboratory (RRID:SCR_018304) is supported in part by the Laura and Isaac Perlmutter Cancer Center Support Grant; NIH/NCI P30CA016087. The RU Center for Clinical and Translational Science Bioinformatics program is supported by the Clinical and Translational Award (CTSA) and the National Center for Advancing Translational Sciences (NCATS), part of the National Institutes of Health. We thank Branka Brukner Dabovic (NYULH – Department of Cell Biology Research) for help with imaging fluorescence samples. We thank the Rockefeller University High-Throughput, Bioimaging and Genomics Resource Centers.

## Material and Methods

### Mice

Male and female *Fah^−/−^* NOD *Rag1^−/−^ Il2rg^null^* (FNRG) mice^24^ were preconditioned with retrorsine^25^ and male *Fah^−/−^ Rag2^−/−^ Il2rg^−/−^* (FRG) mice^26^ with adenoviral urokinase plasminogen activator (uPA) before transplantation with primary human hepatocytes (PHH). NOD *Rag1^−/−^ Il2rg^null^* (NRG) mice were obtained from Jackson Labs (Bar Harbor, ME) and received retrorsine and nitisinone cycling similarly to FNRG mice. Seven week old male thymidine kinase transgenic mice on the NOD SCID *Il2rg^null^* background (TK-NOG)^32^ were obtained from Taconic Biosciences (Germantown, NY) and preconditioned with retrorsine^25^ and 48 hours 0.1mg/ml valganciclovir (Sigma) in drinking water 14 days prior to transplantation. Male transgenic uPA^+/+^/SCID mice were transplanted as previously described.^33^ The following PHH donors were used: cryopreserved or mouse-passaged (mp)PHH1 (HUM4188, Lonza), cryopreserved or mpPHH2 (HUM4129, Lonza), cryopreserved PHH3 (HFCP940, BD Biosciences) and cryopreserved PHH4 (HHF13022, Yecuris). Hepatocytes were genotyped only for the PNPLA3 rs734809 variant. Genotyping was performed by Sanger sequencing using forward 5’-GCCCTGCTCACTTGGAGAAA-3’ and reverse primer 5’-TGAAAGGCAGTGAGGCATGG-3’ as reported.^65^ To make PNPLA3 overexpressing mice, PNPLA3-148I or -148M (both containing the K434E variant^66^) were cloned into SCRPSY to generate VSVg pseudotyped lentiviral particles.^25^ mpPHH1 were isolated from huFNRG, plated in 6 well collagen-coated plates and transduced with lentiviral vectors by 1000g spinoculation. 3 to 5 days after transduction ~5×10^5^ cells/mouse were retransplanted into retrorsine preconditioned FNRG animals as described.^25^ Mice were housed in a 12 hour light/dark cycle at 21°C and 50% humidity. All experiments were conducted under animal use protocols approved by The Rockefeller University, Ghent University and Yecuris.

### Diet challenges

One week prior and during dietary studies, huFNRG, huFRG, NRG and muFNRG mice were changed from oral nitisinone (CuRx, Yecuris cat#20-0028) to intraperitoneal (i.p.) injections at 1mg/kg body weight for three days every 2 weeks. Injection solution was 0.5mg/ml nitisinone dissolved in 500mM NaHCO_3_ pH 7.4 and diluted 1:10 in PBS−/− (Gibco, Thermo Fisher) to final concentration of 50μg/ml. During humanization or ‘murinization’ the FNRG, FRG and TK-NOG lines were maintained on an amoxicillin containing chow diet (modified PicoLab Mouse Diet 20, 5058 - 5B1Q, 0.12% Amoxicillin, irradiated; TestDiet, Richmond, IN) and autoclaved water. One week prior dietary challenge, amoxicillin medicated chow was replaced with regular chow diet (PicoLab Rodent Diet 20, 5053, irradiated; LabDiet, Fort Worth, TX) lacking amoxicillin. For High Fat Diet (HFD) studies, mice were given a diet with 60% kcal fat, 20% carbohydrates and 20% protein (D12492i irradiated; Research Diets, New Brunswick, NJ) and autoclaved water. For Western-style Diet (WD) studies, HFD was combined with 10% w/v sucrose (Fisher Scientific cat#S5-3; Fair Lawn, NJ) in autoclaved water. Sucrose drinking water was freshly made once a week. uPA^+/+^/SCID mice were fed irradiated standard breeding chow (Mouse Breeding complete feed for mice [GE 16.7MJ/kg; 14.0 MJ/kg], Ssniff Spezialdiäten GmbH, Soest, Germany). All diets and drinking water were given *ad libitum*.

### Blood and tissue harvest

Serum was obtained from tail veins, retroorbital or submental venipuncture. Human albumin was quantified as previously described^24^ and serum mouse insulin quantified using Ultra-Sensitive Mouse Insulin ELISA Kit (Crystal Chem cat#90080, Elk Groove Village, IL) following manufacturer’s instructions. ALT and AST activity (Elabscience cat#E-BC-K235 and K236, China) and human ALT protein (Abcam cat#ab234578) were determined following manufacturers instruction, using serum obtained from submental venipuncture 2 to 3 days after last nitisinone injection cycle. Tolerance tests to glucose (GTT), insulin (ITT) and pyruvate (PTT) were performed in FNRG mice in the weeks with no NTBC cycling injections similar as described.^67^ Briefly, age matched mice 4 to 12 weeks after dietary challenge with WD or chow were fasted with free access to water for 6 h prior to GTT and ITT and for 16 h prior to PTT. Approximately 5μl blood was collected from tail bleed before and at indicated intervals for up to 180 min following i.p. injection with 2.5 g/kg body weight glucose (Sigma Aldrich cat#G7021), 1 U/kg body weight human insulin analogue (Eli Lilly, Humulin R U-100), or 2 g/kg body weight pyruvate (Sigma Aldrich cat#P2256), each dissolved in isotonic PBS−/− (Gibco, Thermo Fisher Scientific). Blood glucose concentrations were determined using a GE100 Blood Glucose Monitor (GE, Ontario, CA). Prior to terminal blood and tissue collection (2 to 3 days after last nitisinone injection in *Fah^−/−^* strains), mice were fasted for 6 hours during light cycle with access to water. Blood was collected by submental and/or retroorbital venipuncture and tissue samples were weighed and stored in fixative or frozen on dry-ice and stored at −80°C. Blood was centrifuged with 16,000 x *g* for 15 min at 4°C and plasma was collected and stored at −80°C. Hepatic lipids were extracted from frozen liver tissue as described^67^ and liver and plasma TG, NEFA, total and free cholesterol, and phospholipids were quantified according to manufacturer’s instructions (Wako Diagnostics, Mountain View, CA). Equal volumes of plasma were pooled from three mice and lipoproteins were fractionated by fast protein liquid chromatography (ÄKTA pure FPLC system, GE Healthcare, Pittsburgh, PA, USA) and triglyceride and cholesterol in fractions quantified with reagent kits (Wako Diagnostics, Mountain View, CA).^68, 69^

### Liver histology and quantification

Neutral buffered formalin-fixed (Thermo Fisher Scientific cat#5725) paraffin-embedded liver samples were sectioned at 5μm onto charged slides (Fisher Scientific cat#22-042-924). Slides were dried for 1 hour at 60°C, deparaffinized in xylene, rehydrated through a graded series of ethanol, and rinsed in distilled water for 5 minutes. Sections were then stained with Hematoxylin (Richard-Allan Scientific cat #7211, USA) and Eosin (Leica cat#3801619) using our routine laboratory method described in [Carson, F.L., *Histotechnology: a self instructional text*. 2009, Chicago: ASCP Press., 114-123], and with Picrosirius Red Stain (PSR) (Abcam cat#ab150681) and Masson's Trichrome (Newcomer Supply cat#9179A, USA) in accordance with manufacturer’s supplied procedures. For chromogenic immunohistochemistry staining, unconjugated, polyclonal rabbit anti–Human Nuclear Mitotic Apparatus protein (Abcam Cat# 97585 Lot# GR268490 RRID AB_10680001),^27^ and unconjugated, mouse anti-Human Glutamine Synthetase (Ventana Medical Systems Lot# V0001337 RRID AB_2861318) clone GS-9^70^ were used.^70^ were used. Slides were scanned using a Leica Aperio AT2 System and digitally archived via eSlide Manager. To preserve lipids with and without fluorescent proteins for histopathological analysis, fresh liver samples haven been fixed in 4% paraformaldehyde (Electron Microscopy Sciences) for 18 hours, cryoprotected at 4ºC with 10% and 18% sucrose w/v for 24 hours each before OCT (Sakura Finetek) embedding and storage at −80ºC. OCT-embedded frozen specimens were cryosectioned at 5 μm onto charged slides (Fisher Scientific cat# 22-042-924). Slides were air-dried at room temperature overnight and fixed in 10% neutral buffered formalin for 1 hour at room temperature. Slides were then rinsed in distilled water, stained with BODIPY (Invitrogen cat# D3922) and Oil Red O (Sigma-Aldrich cat# O1391). For BODIPY staining, slides were incubated in Reaction Buffer (Ventana cat# 950-300) for 10 minutes and stained with BODIPY (1:1000 dilution) and DAPI (1:1000 dilution) in PBS for 60 minutes, and covered using ProLong™ Diamond Antifade Mountant (Invitrogen, cat# P36961). BODIPY, RFP, and DAPI were imaged on a Hamamatsu Nanozoomer 2.0 HT and digitally archived via SlidePath. For Oil Red O staining, sections were stained in Oil Red O solution for 10 minutes, rinsed with distilled water for 5 minutes, counterstained with hematoxylin (Richard-Allan Scientific cat# 7211,) for 1 minute and rinsed in distilled water for 1 minute. Samples were blued in 1.2% ammonium hydroxide solution for 1 minute, rinsed in distilled water for 1 minute, and coverslipped using a 50% glycerol solution. Slides were scanned using a Leica Aperio AT2 System and digitally archived via eSlide Manager. For fluorescence image analyses SVS image files derived from Hamamatsu Nanozoomer 2.0 HT were extracted and saved as uncompressed multiple 40x magnification tile images in TIFF format using FIJI (https://imagej.net/Fiji/Downloads)^71^ and NDPITools plugin.^72^ TIFF files of tile images were split into fluorescence channels (DAPI, BODIPY or RFP) and saved as 8-bit grayscale images. Masks were generated around human hepatocytes nuclei approximated by diameter size in DAPI images, enlarged three folds to approximate the size of human hepatocytes in chimeric livers and used for measuring BODIPY and RFP signal intensities. RFP and BODIPY intensities per approximated human hepatocyte were plotted as two dimensional density plots using Python. Histological scores were conducted from H&E stained samples by clinical pathologist (CF and MP), using the NAS system from NASH-CRN.^39^ Steatosis grade (0 :< 5%, 1: 5-33%, 2: >33-66% and 3:> 66%), lobular inflammation (0: no foci, 1: <2 foci / 200x field, 2: 2-4 foci, 3: >4 foci) and ballooning degeneration (0: none, 1: few ballooning cells, 2: many cells / prominent ballooning) were used to generate an overall score (NAS). Fibrosis staging was evaluated separately in PSR stained samples using the same system,^39^ with the following scores 0: none, 1: perisinusoidal or periportal, 1A: mild, zone 3, perisinusoidal, 1B: moderate, zone 3, perisinusoidal, 1C: portal/periportal.

### Liver tissue transcriptional analyses

Around 25 mg of freshly harvested liver tissue was placed into 1ml 4ºC TRIzol (Life Technologies, USA) with 500μl of 1mm glass beads (BioSpec Products cat# 11079110) and homogenized using MagNA lyser (Roche, USA), weighed and stored at −80ºC. For RNA extraction, 500μl of liver/Trizol mixture was used for chloroform extraction using MaXtract™ (Qiagen, cat# 129056) and RNeasy® Mini Kit (Qiagen, cat# 74104) coupled with on-column DNaseI treatment (Qiagen cat#79254) following manufacturer’s instructions. Extracted RNA was stored at −80ºC. For qRT-PCR, cDNA was generated from liver RNA using SuperScript IV VILO (Thermo Fisher Scientific cat#11766050) following manufacturer’s instructions. qRT-PCR was performed in 96-well Human Fatty Liver Array plates (Thermo Fisher Scientific cat # 4391524) on a QuantStudio 3 System (Applied Biosystem) for the following genes: *ABCA1, ACACA, ACADL, ACLY, ACOX1, ACSL5, ACSM3, ADIPOR1, ADIPOR2, AKT1, APOA1, APOB, APOC3, APOE, ATP5C1, CASP3, CD36, CEBPB;CEBPB-AS1, CNBP, CPT1A, CPT2, CYP2E1, CYP7A1, DGAT2, FABP1, FABP3, FABP5, FAS, FASN, FOXA2, FOXO1, G6PC, G6PD, GCK, GK, GSK3B, HMGCR, HNF4A, IGF1, IGFBP1, IL1B, INSR, IRS1, LDLR, LEPR, LPL, MAPK1, MAPK8, MLXIPL, MTOR, NDUFB6, NFKB1, NR1H2, NR1H3, NR1H4, PCK2, PDK4, PIK3CA, PIK3R1, PKLR, PNPLA3, PPA1, PPARA, PPARG, PPARGC1A, PRKAA1, PTPN1, RBP4, RXRA, SCD, SERPINE1, SLC27A5, SLC2A1, SLC2A2, SLC2A4, SOCS3, SREBF1, SREBF2, STAT3, XBP1*. Transcriptional changes were calculated over mean of housekeeping genes: *GAPDH, HPRT1, GUSB, ACTB, B2M, HMBS, IPO8, PGK1, RPLP0, TBP, TFRC*. Data were analyzed and graphed in R (version 4.0.2). For bulk RNA sequencing (RNAseq) the quantity and integrity of extracted liver RNA were assessed using an Agilent 2100 Bioanalyzer (Agilent Technologies, Palo Alto, CA). Library construction and sequencing were performed by Novogene USA (Sacramento, CA), using Illumina S4 flowcell on Novaseq 6000 platform with a paired-end read length of 150 bp. Downstream RNAseq analysis was performed using a combination of programs. Alignments were parsed using STAR and differential expression was determined through limma/voom.^73^Alignments were parsed using STAR and differential expression was determined through limma/voom.^73^ GO, KEGG, REAC and TF enrichment were determined with the g:profiler R package.^74^ Reference genome and gene model annotation files were downloaded from genome website browser (NCBI/UCSC/Ensembl) directly. Indices of the reference genome were built using STAR^75^ and paired-end clean reads were aligned to human or mouse reference genome using STAR (v2.5), setting the outFilterMismatchNmax argument to 2, from which gene level count tables were built using HTseq-count.^76^ Gene-level read counts were then processed using the limma suite of tools^73^ first with a voom with quality weights transformation, followed by linear model fitting to determine differentially expressed genes. The P values for individual genes were adjusted using the Benjamini & Hochberg method. Corrected P-value of 0.05 and absolute foldchange of 1 were set as the threshold for significantly differential expression. Geneset testing on the entire dataset was performed using the camera() function to determine geneset direction using genesets extracted from MsigDB.^77^ Select genesets were plotted as dotplots from the camera output. Gene Ontology (GO), Kyoto Encylopedia of Genes and Genomes (KEGG), and REACTOME database enrichment analysis of differentially expressed genes was further implemented with the g:profiler R package,^74^ in which gene length bias was corrected. Terms with corrected P-values less than 0.05 were considered significant. The high-throughput sequencing data from this study have been submitted to the NCBI Sequence Read Archive (SRA) under accession number ####.

### Data plotting and statistical analysis software

Unless stated otherwise, statistical analyses were performed in GraphPad Prism 9 (San Diego, CA). Statistics are indicated in graphs if p<0.05.

## Author contributions

MK, EM, SS, BE, CMR and YPJ designed research; MK, EM, SS, CGF, JML, JLP, MT, BR, IRL, C.Z., BZ, AFS, CQ, LF, AWA, WMS, SB, GL, LC and YPJ performed experiments; MK, EM, SS, CGF, JML, JLP, MT, BR, IRL, CZ, BZ, AFS, CQ, LF, AWA, YL, DEC, MP, LD, MG, PM, BAE, CMR and YPJ analyzed data; HS, NDT and RC provided new reagents/analytic tools; DEC, PM, CMR and YPJ secured funding; MK, EM, and YPJ wrote the paper; YPJ directed the study.

