## Supplemental Figures for "Human hepatocyte PNPLA3 148M exacerbates rapid non-alcoholic steatohepatitis development in chimeric mice"

### Supplemental Figure S1: Human hepatocyte steatosis in chimeric models

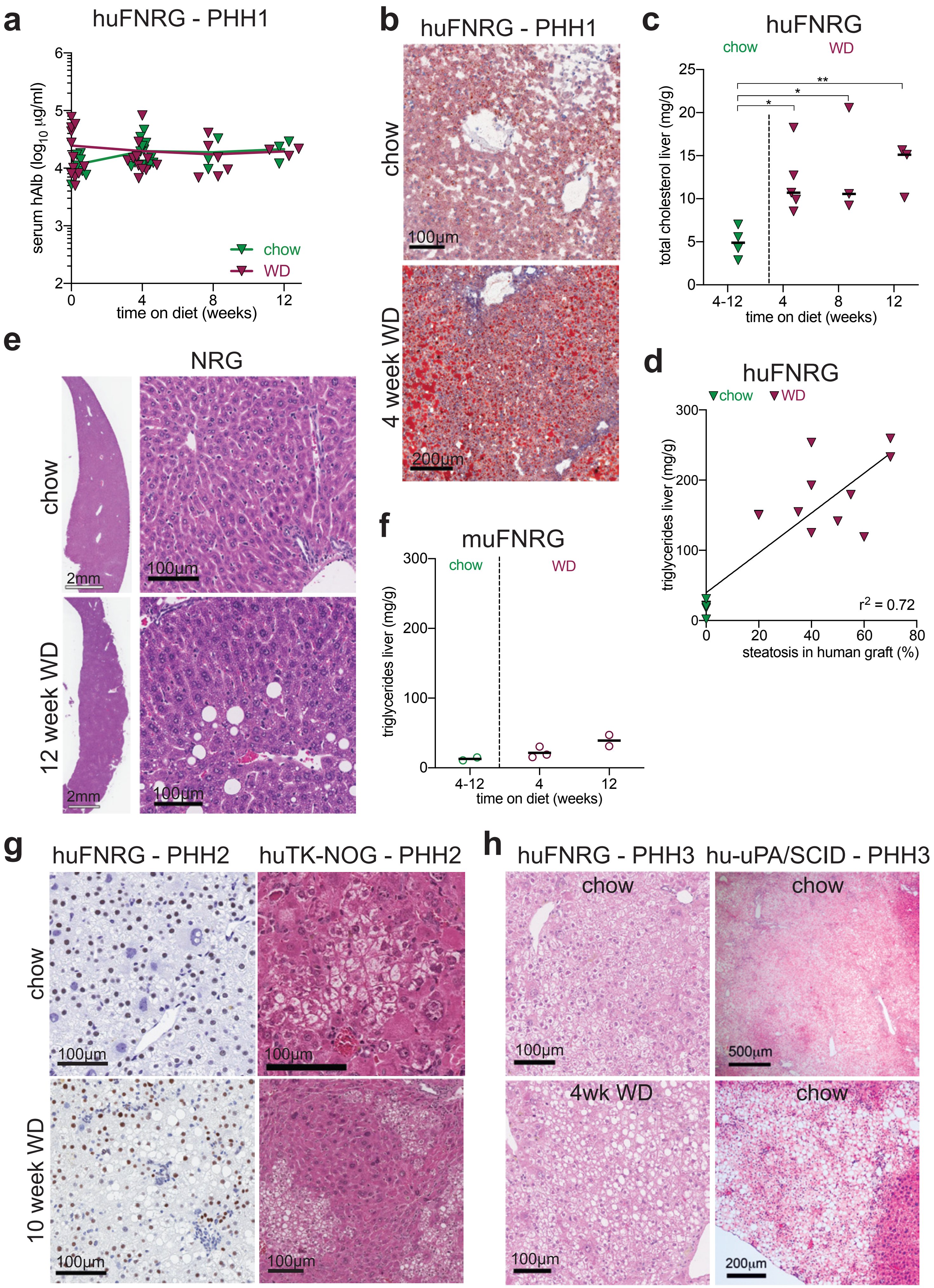

### Supplemental Figure S2: Systemic metabolic effects of WD feeding

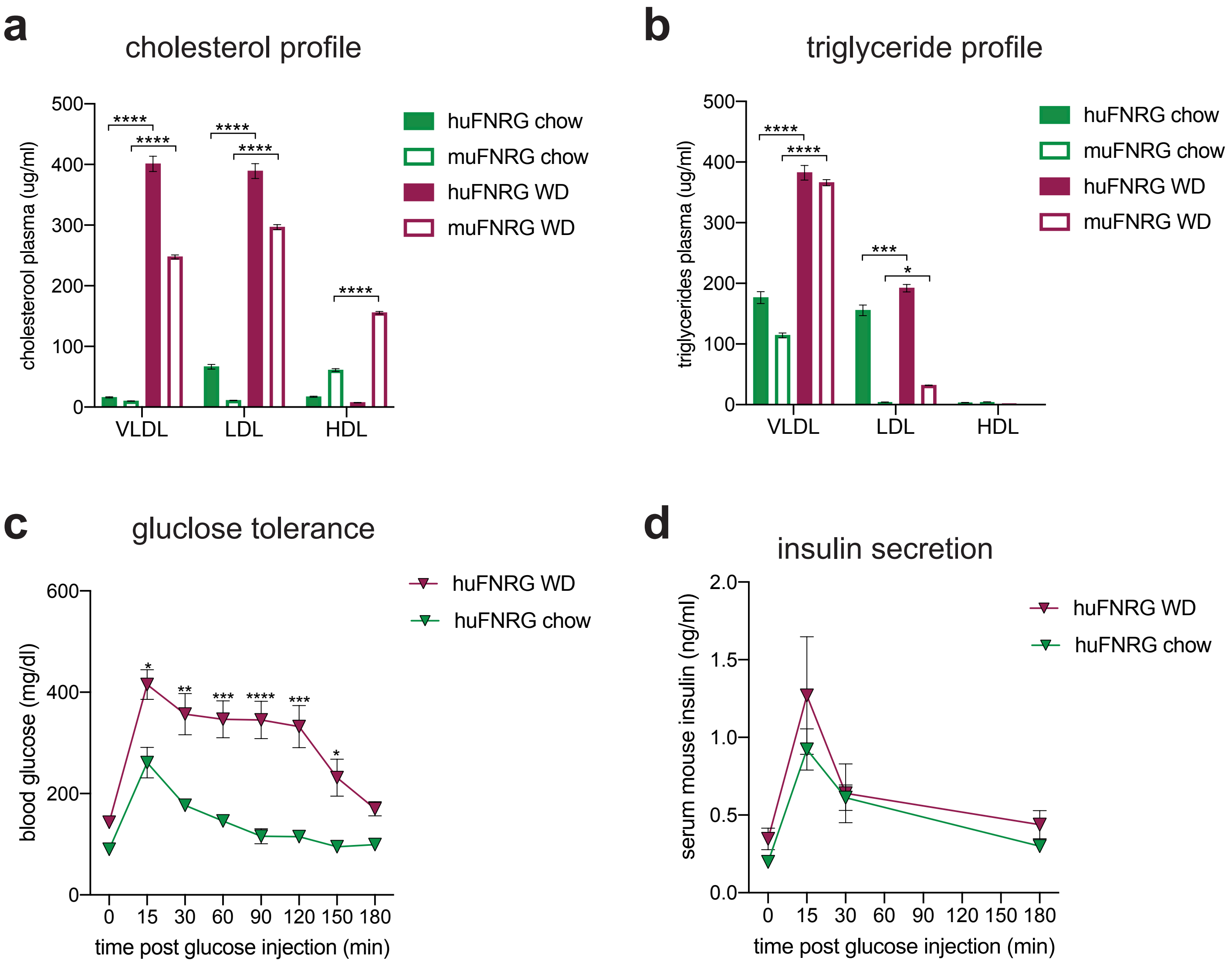

Supplemental Figure S3: Mild steatohepatitis in huFNRG mice on WD

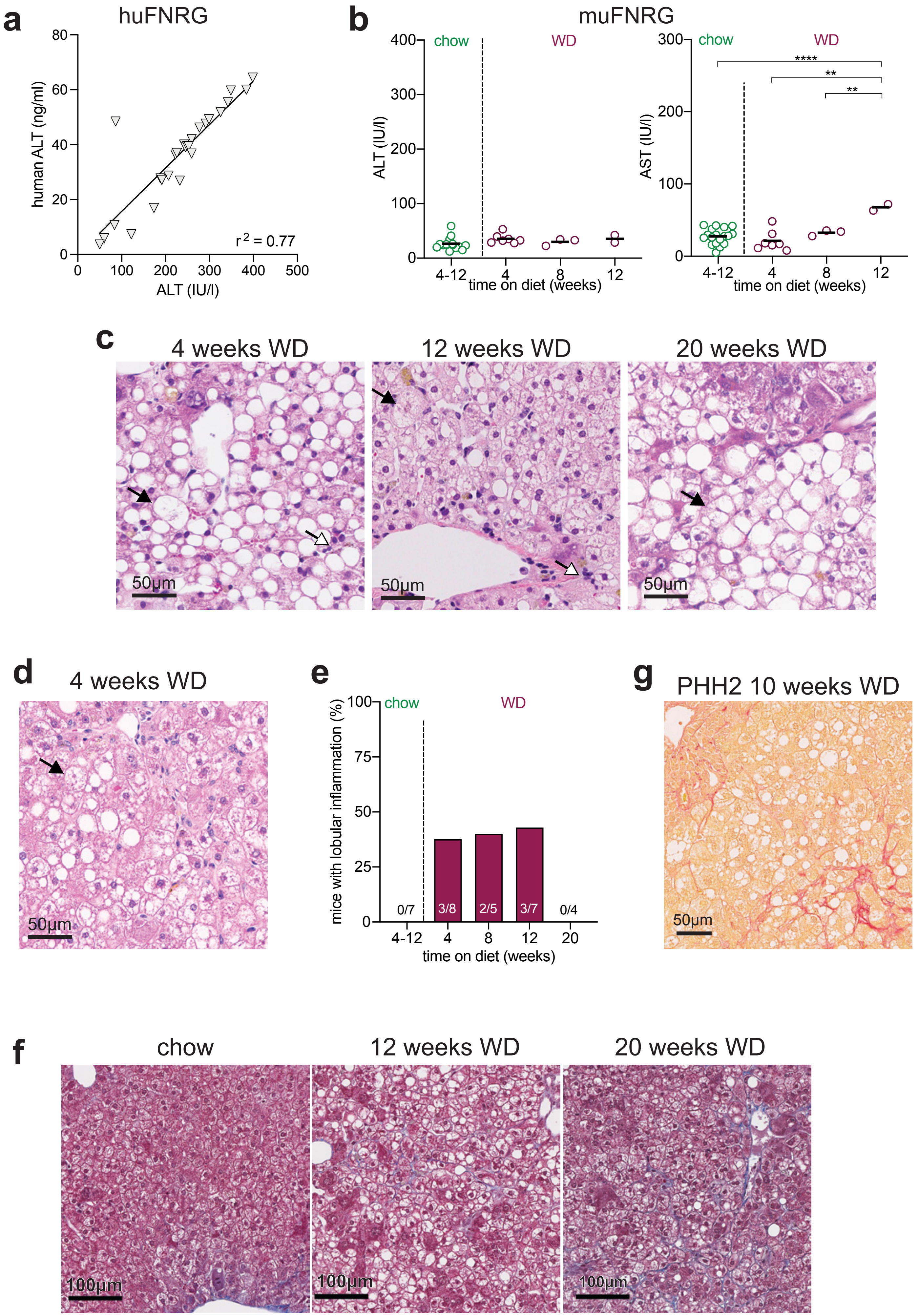

### Supplemental Figure S4: Transcriptional changes in huFNRG mice on Western Diet

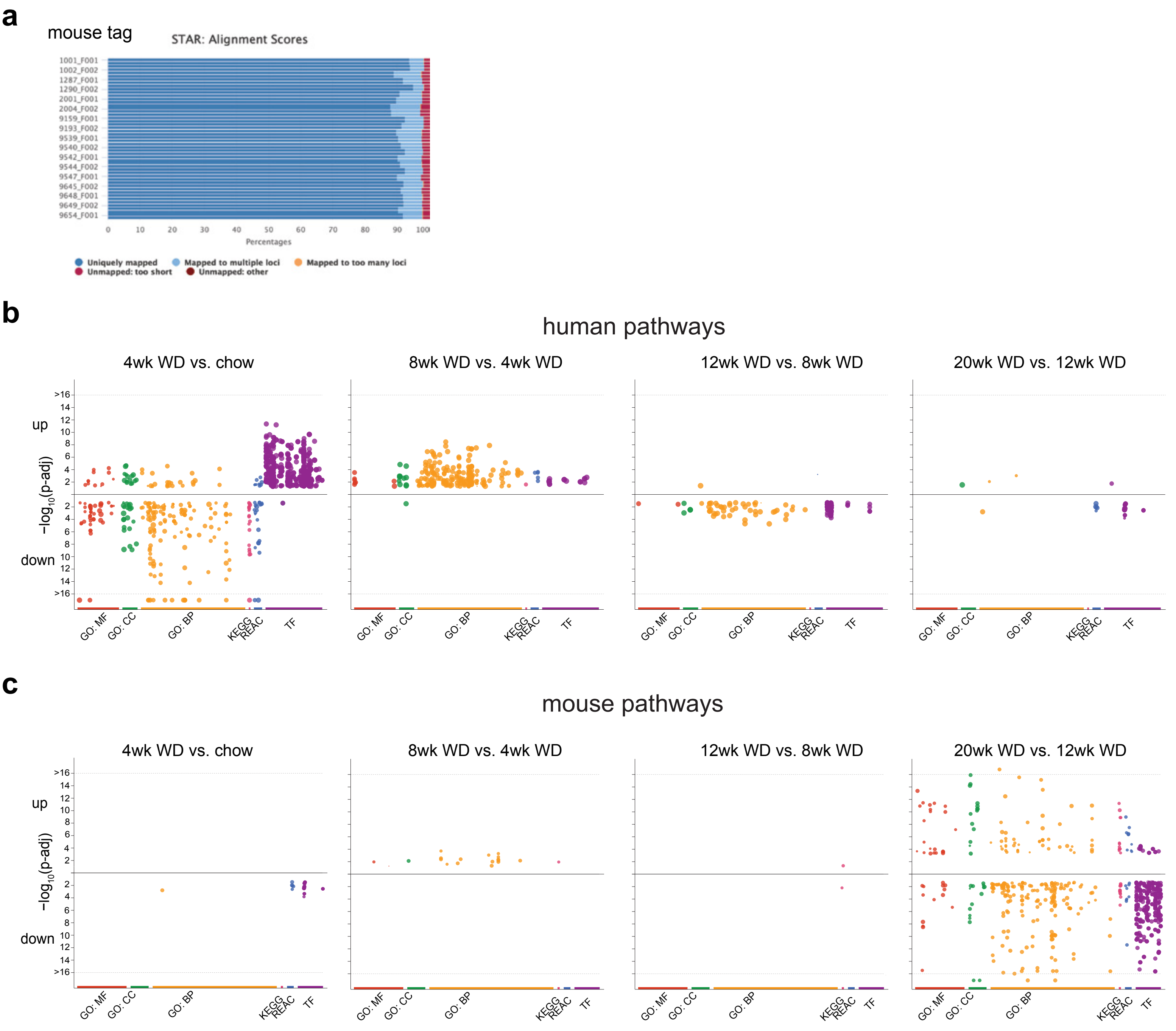

### Supplemental Figure S5: 148M-huFRG mice develop steatohepatitis

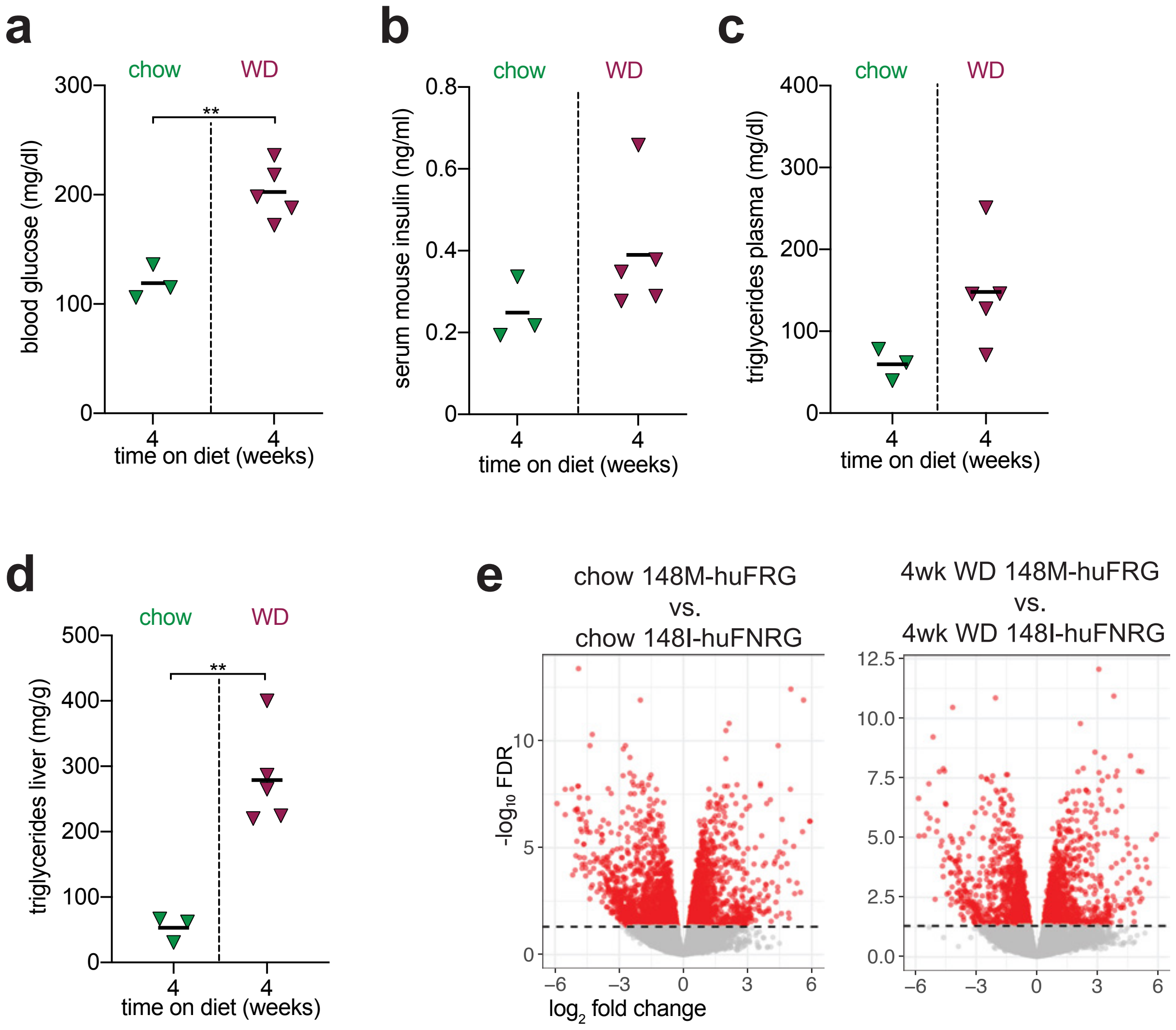

### Supplemental Figure S6: PNPLA3 transgenic huFNRG mice

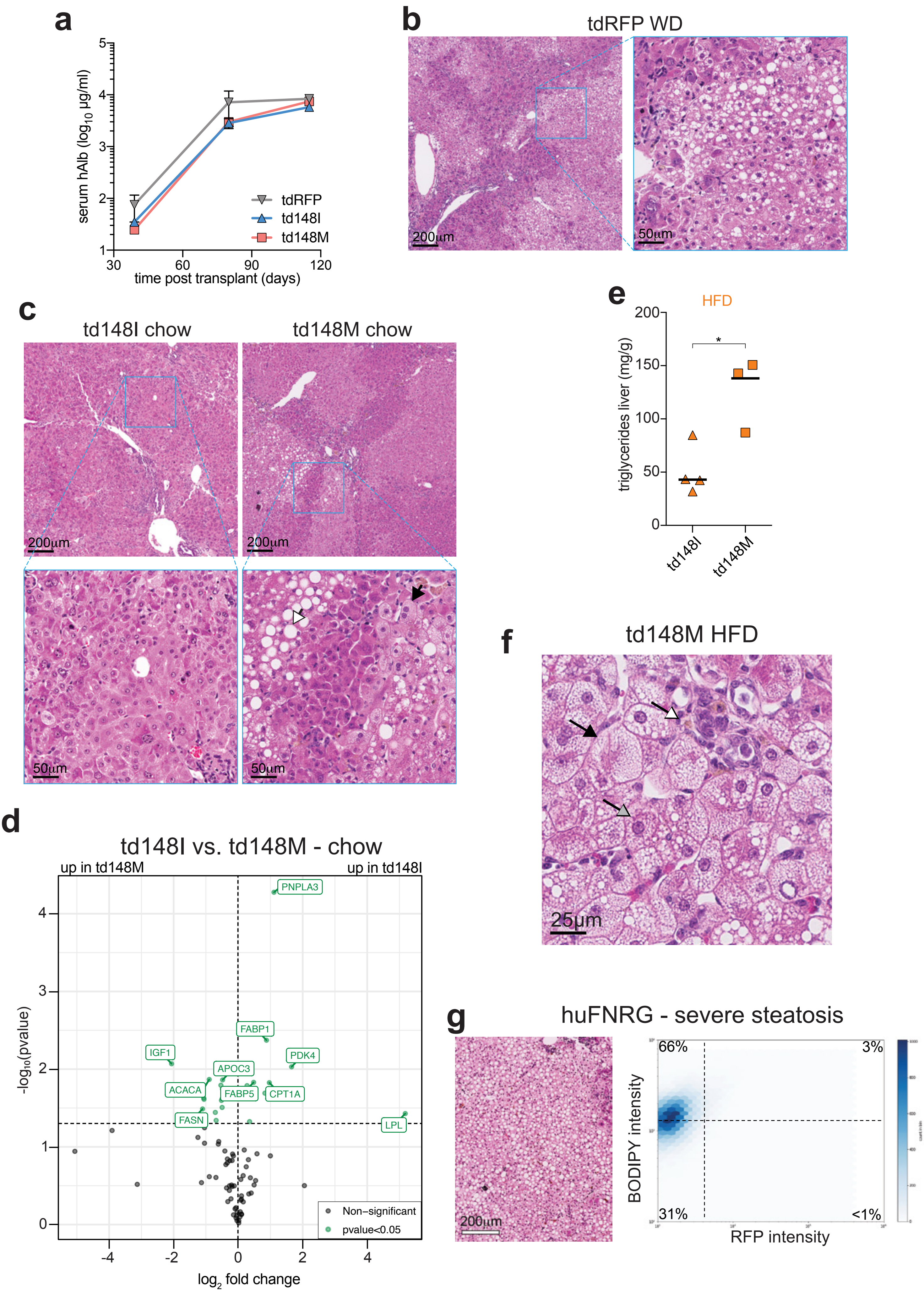
